# Developmental neuroanatomy of the Rosy Bitterling *Rhodeus ocellatus* (Teleostei: Cypriniformes)—A microCT study

**DOI:** 10.1101/2021.10.08.463635

**Authors:** Wenjing Yi, Thomas Mueller, Martin Rücklin, Michael K. Richardson

## Abstract

Bitterlings are a group of teleost fish (Cyprinifromes: Acheilanathidae) notable for their brood parasitic lifestyle. Bitterling embryos develop as parasites inside the gill chamber of their freshwater mussel hosts. However, little is known about brain development in this species. Here, we have imaged the development of the brain of the Rosy Bitterling (*Rhodeus ocellatus*) at four embryonic stages (165, 185, 210, 235 hours post-fertilization) using micro-computed tomography (microCT) with special emphasis on developmental regionalization and brain ventricular organization. We provide a detailed neuroanatomical account of the development of the brain divisions with reference to The *Atlas of Early zebrafish Brain Development* and the updated prosomeric model. Segmentation and three-dimensional visualization of the ventricular system were performed in order to clarify changes in the longitudinal brain axis as a result of cephalic flexure during development. During early embryonic and larval development, we find that histological differentiation, tissue boundaries, periventricular proliferation zones, and ventricular spaces are all recognizable using microCT. Importantly, our approach is validated by the fact that the profile of CT values displayed here in the bitterling brain are consistent with genoarchitecture identified in previous studies. We also find developmental heterochrony of the inferior lobe in the Rosy Bitterling compared to the zebrafish. Our study provides a foundation for future studies of the brain development in the Rosy Bitterling, a valuable model species for studying the evolutionary adaptations associated with brood parasitism.

## 1. INTRODUCTION

Bitterlings, a group of freshwater teleosts, are valuable model species in behavioral, population and evolutionary ecology due to their brood parasitic life history. Their peculiar life style involves the laying of eggs by the bitterling in a host mussel, a phenomenon that has been recognized for more than a century (Boeseman et al., 1938; Chang, 1948; Duyvené de Wit, 1955; Kitamura et al., 2012; Mills and Reynolds, 2003; Olt, 1893; Reichard et al., 2007; Rouchet et al., 2017; Smith, 2016; Wiepkema, 1962). Noll (1877) was the first to show that the embryos of European bitterling (*Rhodeus amarus*) develop in the gill chamber of their host mussel. This location provides a sheltered environment which may protect the developing embryos from predators (Aldridge, 1999; Liu et al., 2006; Reichard et al., 2007; Smith et al., 2004). In a recent study (Yi et al., 2021) we compared developmental sequences of the Rosy Bitterling (*Rhodeus ocellatus*) to the zebrafish (*Danio rerio*); the latter is a non-parasitic teleost which lays its eggs into the open water (Kimmel et al., 1995; Lawrence, 2007). Our study identified a number of heterochronic (timing) shifts, including the relative pre-displacement of hatching, and the relative delay of development of the pectoral fins, in the bitterling.

The specialized development of the bitterling, in relation to its brood parasitic lifestyle, make it a potentially interesting model for the study of developmental mechanisms underlying brain evolution. In this study, we generated an atlas of the developing bitterling brain as a reference for cross-species comparisons. To this end, we compared bitterling brain development with published data for the zebrafish. The comparison to the zebrafish is useful for two reasons. First, the zebrafish is firmly established as a genetic model system that has been thoroughly investigated with regard to its embryonic, postembryonic, and larval development (Mueller and Wullimann, 2003; Mueller and Wullimann, 2016; Mueller et al., 2006; Wullimann, 2009; Wullimann and Knipp, 2000; Wullimann and Mueller, 2004). Second, both, the bitterling and the zebrafish belong to the group of carp-like (cyprinid) teleosts and have similar adult brain anatomy. However, as we show, these two species show distinct developmental timing differences (heterochronies). The developmental staging atlas of the bitterling brain, that we describe here, provides a foundation for future studies on the molecular mechanisms underlying these heterochronies.

In the vertebrates, the central nervous system, including its anterior part, the brain, develops from the neural tube (Richardson and Wright, 2003; Schmitz et al., 1993; Wullimann, 2009). In teleosts, including bitterlings, the development of the neural tube involves secondary neurulation (Schmitz et al., 1993). The lumen of the definitive central nervous system contains cerebrospinal fluid (Lowery and Sive, 2009). The spinal cord has a narrow lumen, the central canal. The brain, by contrast, has an inflated, irregular lumen which form a series of brain ‘ventricles’ (Korzh, 2018). The four main divisions of the vertebrate are: (i) the secondary prosencephalon; (ii) the diencephalon; (iii) the mesencephalon and (iv) the rhombencephalon. Accordingly, the brain ventricular system has been divided into the prosencephalic ventricle (including the telencephalic ventricle and the hypothalamic part of the classic diencephalic ventricle; Nieuwenhuys and Puelles, 2016); the 3^rd^ ventricle (the lumen of diencephalon proper); the mesencephalic ventricle; and the 4^th^ ventricle (the lumen of the rhombencephalon).

In this study we use the updated prosomeric model of Puelles and Rubenstein (2003) for dividing neuromeres in the secondary prosencephalon as well as for defining other brain divisions. The prosomeric model recognizes longitudinal zones and transverse zones (corresponding to the neuromeres) is a powerful paradigm for vertebrate cross-species comparisons. The model is based on the conserved expression patterns of developmental genes and on morphological characters. These data permit the demarcation of the CNS into developmental units (morphogenetic entities) along the neuraxis of a range of vertebrate species. In teleosts, patters of cell migration and differentiation also identify prosomeric units and can be used to map them (Mueller and Wullimann, 2003; Mueller and Wullimann, 2016; Mueller et al., 2006; Wullimann, 2009; Wullimann and Knipp, 2000; Wullimann and Mueller, 2004).

The prosomeric model analyzes brain regions along the general brain axis and according to the mediolateral extent of brain regions. Longitudinally, the neural tube has four compartments divided by the *sulcus limitans* of Wilhelm His, Sr. (His, 1895); reviewed by Puelles (2019). The four compartments are: the roof plate dorsally, the alar plate dorsolaterally, the basal plate ventrolaterally, and the floor plate ventrally.

According to Nieuwenhuys and Puelles (2016), the rhombencephalon consists of twelve neuromeres (rhombomeres) whereby the isthmus is designated r0, and the remaining rhombomeres r1–r11) in longitudinal series. In cyprinids such as the Goldfish (*Carassius auratus*), rhombomeres r2–r6 correspond to the mammalian pons, whereas the rhombomeres 7-8 correspond to the mammalian medulla oblongata (Gilland et al., 2014; Rahmat and Gilland, 2019). Rhombomeres r2–r6 form clearly segmented neural clusters while rhombomeres r7–r8 are only poorly delimited (Ma et al., 2009).

In the zebrafish, r8 is twice as large as the rostral rhombomeres; the latter are composed by multiple crypto- or pseudo-rhombomeres that can only be clearly by molecular markers, as is the case in their avian homologs (Cambronero and Puelles, 2000). The rostral subdivision of neuromeres is more complex, because bending of the neural tube (the cephalic flexure, Hauptmann and Gerster, 2000; Mueller and Wullimann, 2009) has obscured the delineation of the longitudinal neuraxis (Mueller and Wullimann, 2009; Puelles, 2019; Vernier, 2017; Wullimann and Rink, 2002).

Here, we visualize the development of the Rosy Bitterling (*Rhodeus ocellatus*) brain using micro-computed tomography (microCT, x-ray microscopy or µCT) with a specific focus on developmental regionalization and brain ventricular organization. MicroCT is a high-resolution, non-destructive, three-dimensional (3D) imaging technique (Babaei et al., 2016; Metscher, 2009a) that has recently begun to be used in developmental studies (Wong et al., 2015). The contrast of CT scans is based on the absorption of x-ray radiation passed through the sample. Highly mineralized structures such as bones and teeth have a higher attenuation coefficient (CT value; Hounsfield units). This makes them appear brighter in the final image, and easier to recognize than soft tissues. For the visualization of the latter, a treatment with a contrast agent is needed. In this work, we used phosphotungstic acid (Metscher, 2009b), which enhances the visualization of soft tissues such as muscles, nerves, and blood vessels.

In this study, we extend MicroCT techniques to the developing bitterling brain. Our goal is to establish microCT and 3D visualization as complementary methods for cross-species comparisons of the morphology of both the developing and the mature brain. Our results indicate that microCT is a valuable modality for the study of the ventricles, and for discriminating white matter from proliferative and postmitotic cell masses.

## 2. MATERIALS AND METHODS

### Sample preparation

Embryos of six developmental stages (135, 150, 165, 185, 210, 235 hours post fertilization (hpf); Table 1) were obtained by *in vitro* fertilization following the method of Nagata and Miyabe (1978). Embryos were staged according to Yi et al. (2021). Embryos were incubated at 22.5 ± 1 °C and fixed in 3% buffered paraformaldehyde (pFA) and 1% glutaraldehyde (GA). Digital microphotographs of fixed samples were obtained with a CCD (charge-coupled device) camera connected to stereo microscope (Nikon SMZ1500). For x-ray contrast enhancement, embryos were stained for at least 24 h in 0.3% phosphotungstic acid (PTA) dissolved in 70% ethanol. Samples were then brought back to 70% ethanol without PTA and mounted in low-melting point agarose for non-shift scanning in Gilson® Pipetman® tips.

**Table 1.**
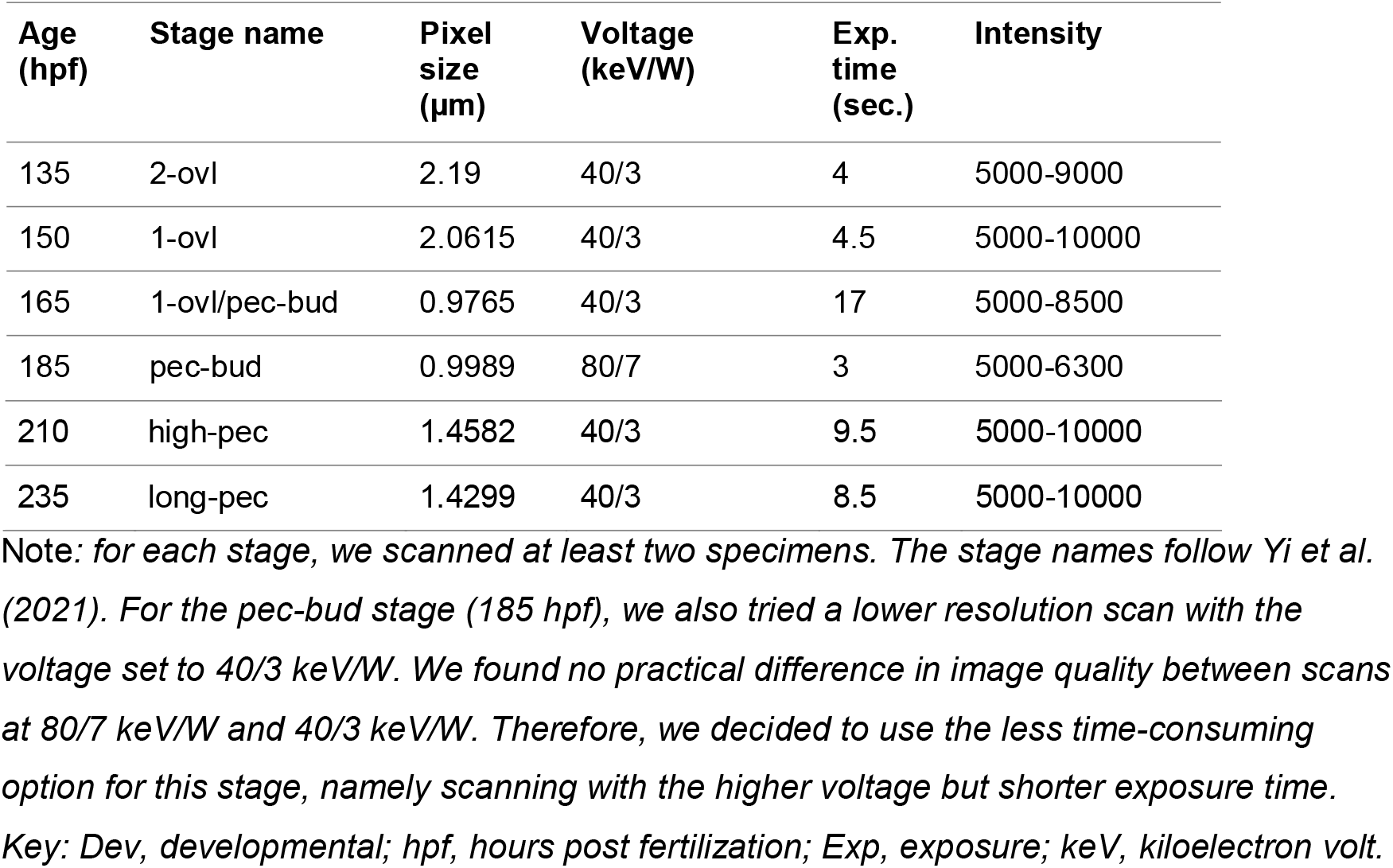
Sample information and microCT scanning parameters.

### MicroCT scanning

Attenuation-based microtomographic images were acquired using an Xradia 520 Versa 3D X-ray microscope (Zeiss), with the x-ray tube voltages source set at 80/7 or 40/3 keV/W (keV: kiloelectron volts). A thin LE1 filter was used to avoid beam-hardening artifacts. During the CT scanning, the sample was placed on a rotation table and projection images were acquired over an angular range of 180 degrees. To obtain high resolution images, a 4x CCD (charge-coupled device) optical objective was used. Images were acquired with voxel sizes (volumetric 3D pixels after reconstruction) of 1-1.5 µm, and tomographic reconstructions were made with the resident software (XMReconstructor). Reconstructed images were exported as TIFF files and loaded into Avizo version 9.5 (Thermo Fisher Scientific) for 3D visualization.

### 3D visualization

The reconstructed volume was viewed slice-by-slice as virtual sections using the *Slice* module in the Avizo software (Version: 9.5; Thermo Fisher Scientific). A computational module *Resample transformed image* was applied to register images in the orthogonal direction of the anatomical axis. *Ortho View* was used for interactive orthogonal views in the *xy*, *yz* and *xz* axes simultaneously. The *Volume endering* module was used for 3D viewing. Colormap and opacity were optimized in the rendering settings. Scalebars were added using the *Scalebars* module.

### Annotations on 2D slices

To generate a developmental atlas, serial virtual transverse sections were taken in rostrocaudal sequence from the olfactory bulb to the medulla oblongata. The section plane was parallel to the deep ventricular sulcus (the anterior intraencephalic sulcus or AIS) between the telencephalon and diencephalon. To keep a consistent prosomeric axis annotation, we visualized the transverse sections from the rostral to caudal, the coronals section from dorsal to ventral, and the sagittal sections from medial to lateral along the primary body axis (Figure 1f). Labels were added to each section in using Adobe InDesign® software (Version: 15.0.2; Adobe Systems Inc., San José, California). For anatomical terms see the list of abbreviations (Figure xx). We mainly used the anatomical terms in the *Atlas of Early zebrafish Brain Development* (Mueller and Wullimann, 2005, 2016).

**Figure 1.**
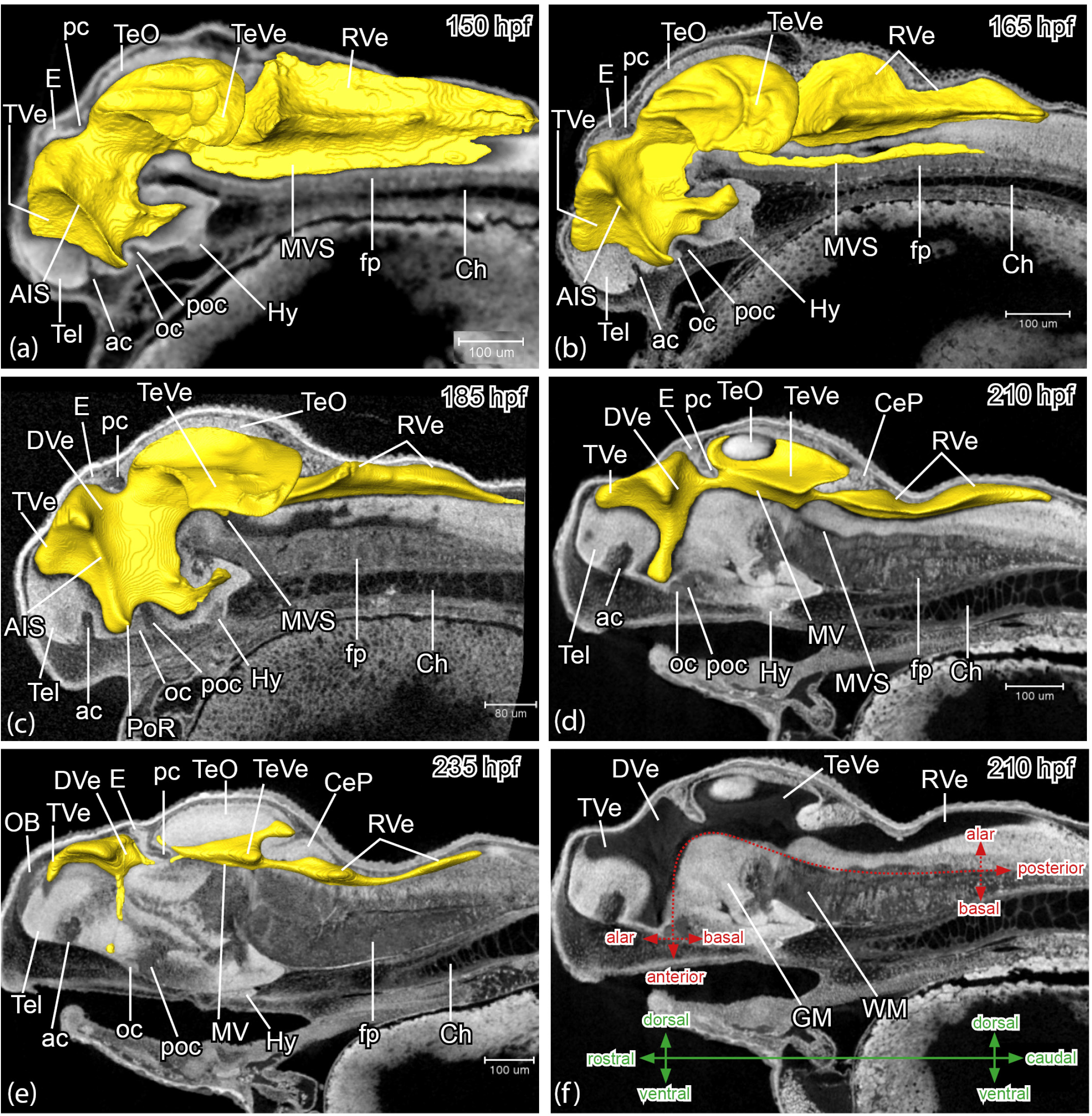
*Rhodeus ocellatus*, development of the brain ventricular system, microCT images, dorsal to the top, head to the left. (a-e) virtual midsagittal sections overlayed with surface view of manually-segmented brain ventricles, and showing the distinct ridge of the anterior intraencephalic sulcus (AIS) and progressive compartmentalization of the ventricular system (a-e). (f) virtual midsagittal section illustrating the black lumen of the brain ventricle, a bright white periventricular layer of gray matter (GM), and a gray peripheral layer of white matter (WM). The red dotted line indicates the neuraxis, with anterior, posterior, alar and basal topological direction for the curved axis. The green axes refer to the linear axial system apply to the brain parts with the terms rostral, caudal, dorsal, and ventral. The (a) stage 1-ovl, 150 hpf. (b) stage 1-ovl/pec-bud, 165 hpf. (c) stage pec-bud, 185 hpf. (d, f) stage high-pec, 210 hpf. (e) stage long-pec, 235 hpf. For annotations, see List of Abbreviations. Scale bars = 100 μm (a, b, d, e. f); 80□μm (c).

### Segmentation of brain ventricles

Segmentation of the brain ventricles was conducted in Avizo in two steps. First, a rough segmentation based on greyscale threshold was achieved semi-automatically, and then polished using *Smooth labels* and *Remove islands* filters. These segmentation results were checked slice-by-slice and were manually corrected. The *Generate Surface* module was used to extract surfaces from the segmentation results. Brain ventricles were colored in yellow using the *Surface view* module, while the rest of the cranial tissues were rendered semi-transparent using the *Volume rendering* module. The segmented model of the brain ventricles was captured in dorsal, ventral, lateral, and rostral views and saved in TIFF file formats. They were annotated using Adobe InDesign®.

## 3. RESULTS

We have studied brains of the Rosy Bitterling (*Rhodeus ocellatus*) using microCT. The developmental ages were 165, 185, 210 and 235 hpf (hours post fertilization). Throughout this study we have used the stage table of development of the Rosy Bitterling generated by Yi et al. (2021). To avoid confusion regarding the anatomical orientation, we follow Herget et al., (2014) and use the terms rostral, caudal, dorsal and ventral as in classical descriptions for the primary embryonic axis. To take account of the curvature of the neuraxis, we sometimes use the terms anterior, posterior, alar and basal as alternatives (Figure 1f). Our results are divided into two sections. First, we describe the development of the ventricular system. Second, we present a developmental atlas of the bitterling brain.

### A. Development of the brain ventricular system of the Rosy Bitterling

#### General description

The greyscale values observed on virtual microCT sections are influenced by PTA staining, tissue density, and cell type. In the brain, the gray matter generally appeared brighter than the white matter (for example, compare the telencephalon and anterior commissure in Figure 1, Tel and ac). The white matter with its myelinated axons and tracts yielded greyscale values typical of low-density soft tissues (Figure 1, ac, poc, pc). In contrast, the brain ventricle, essentially a hollow space filled with cerebrospinal fluid (CSF), showed a lower density than the nervous tissue of the brain. It was visible in CT scans as the darkest part (Figure 1a-f, rendered in yellow color in Figure 1a-e). Therefore, the radiological appearance of the brain encompasses, broadly speaking, three greyscale values: (i) a black lumen: the ventricular system; (ii) a bright white periventricular layer of gray matter, and (iii) a gray-coloured peripheral layer of white matter (Figure 1f).

The forebrain ventricle is divided dorsally by the anterior intraencephalic sulcus (AIS) into the anterior telencephalic ventricle (TVe) and the posterior diencephalic ventricle proper (DVe or the 3^rd^ ventricle; Figure 1). During the eversion of the telencephalic ventricle, the dorsal part of the AIS shows concomitant enlargement (Figure 1). Around the preoptic recess (PoR), the optic recess region (ORR) was recognizable, bordered by the anterior commissure (ac) and postoptic (poc) commissures (see locations of ac, poc and PoR in Figure 1c). The hypothalamic ventricle topographically locates caudal (topologically basal) to the optic recess region, and topologically anterior to the 3^rd^ ventricle, including the lateral recess (LR) and the posterior recess (PR; Figure 2).

**Figure 2.**
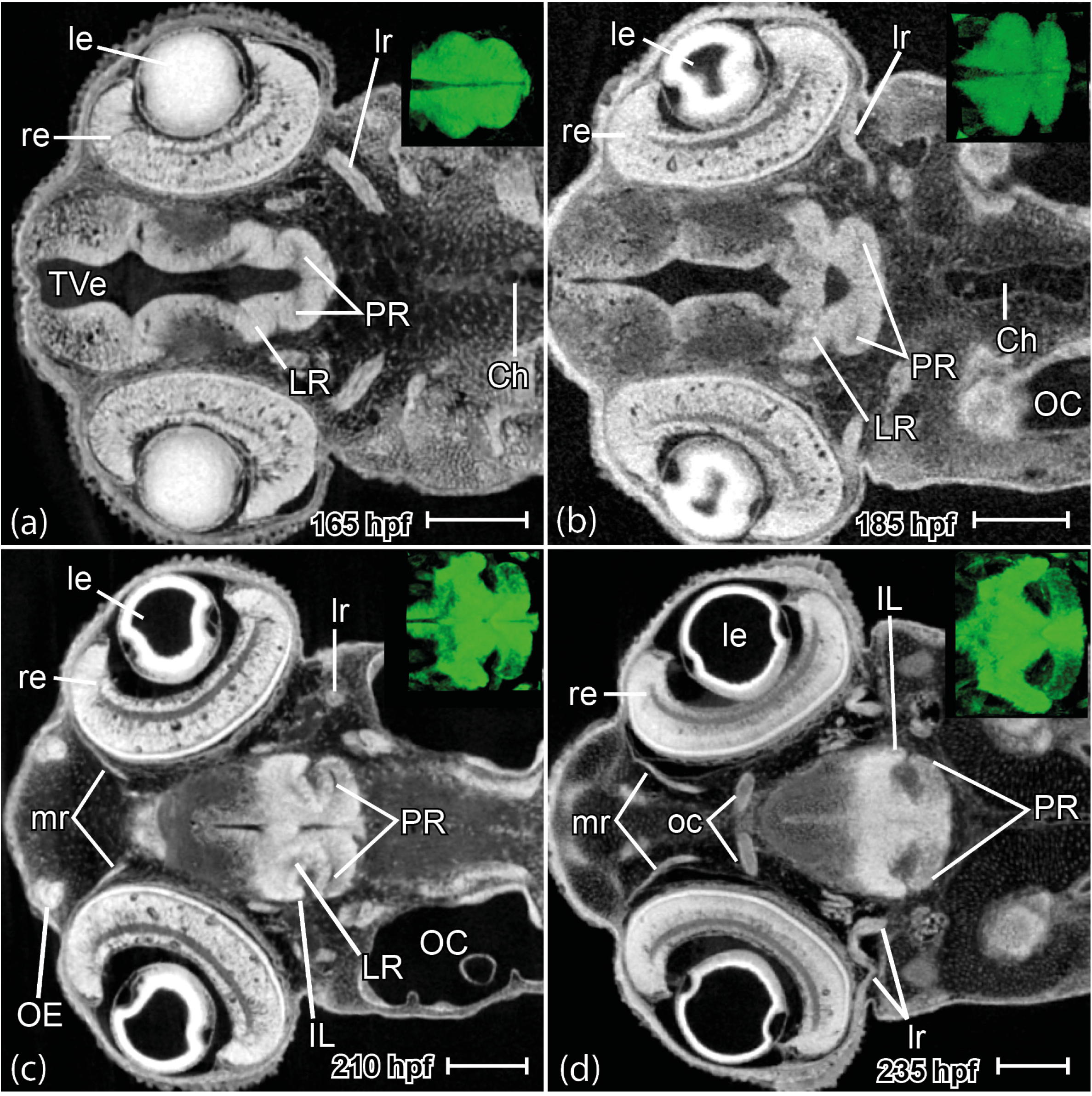
*Rhodeus ocellatus*, development of the lateral recess and posterior recess in the hypothalamus region. Virtual coronal sections, head to the left. At the right upper corner is the 3D volume rendering of the hypothalamus region seen from the dorsal side. (a) stage 1-ovl/pec-bud, 165 hpf. (b) stage pec-bud, 185 hpf. (c) stage high-pec, 210 hpf. (d) stage long-pec, 235 hpf. For annotations, see List of Abbreviations. Scale bars = 100 μm

The mesencephalic ventricle encompasses the paired tectal ventricle (TeVe), the median ventricle (MV), and the median ventricular sulcus (MVS) (Puelles, 2019). The TeVe projects dorsolaterally from the median ventricle (Figure 3a), covered by the subarea of the alar plate of the midbrain (optic tectum). A transient median ventricular sulcus (MVS) branches off from the ventral bottom of the median ventricle and shows separately at the floor plate area (Figure 3b, inset in the upper right corner). The median ventricle extends caudally to the rhombencephalic ventricle (RVe or the 4^th^ ventricle; Figure 3; García-Lecea et al., 2017; Korzh, 2018).

**Figure 3.**
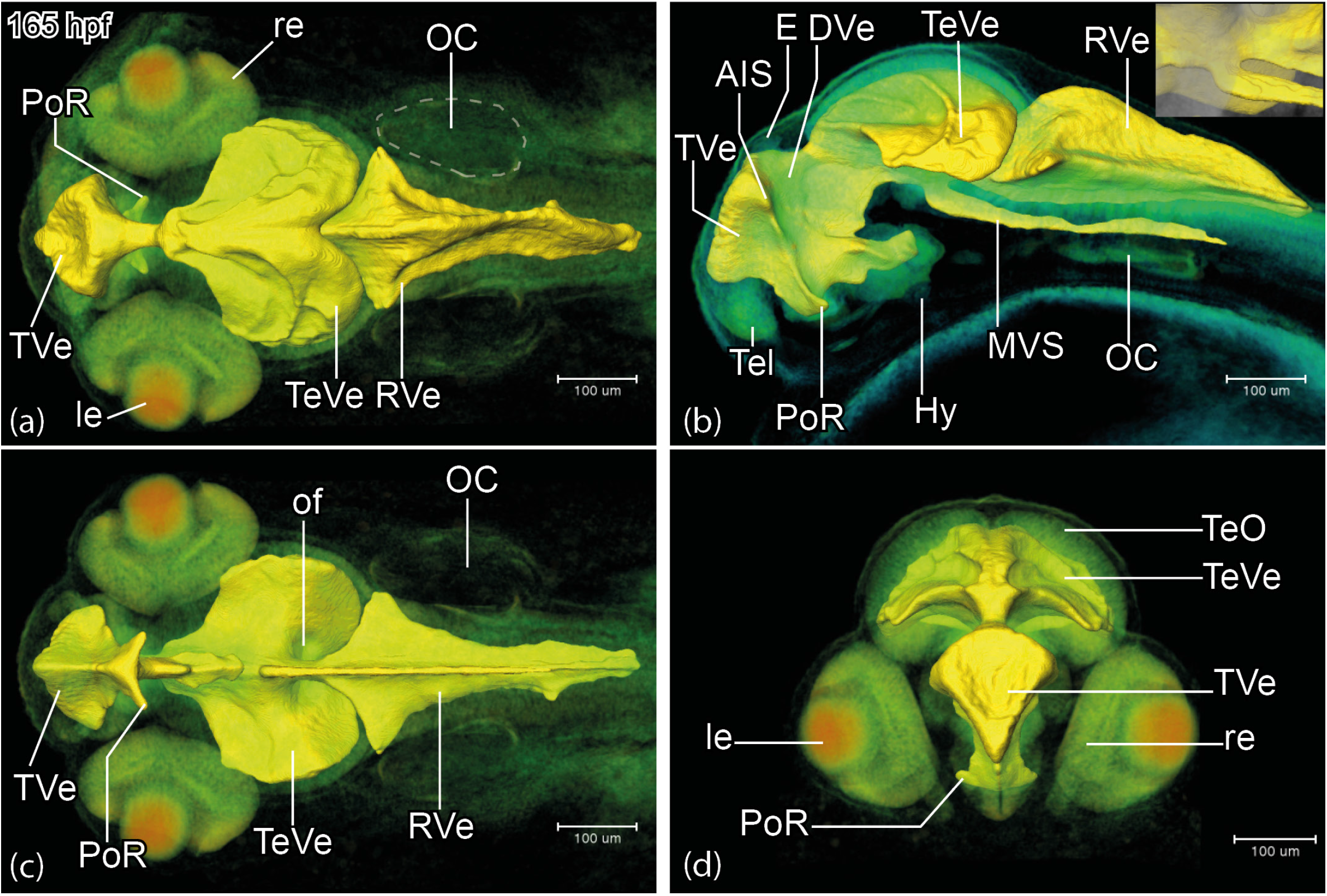
*Rhodeus ocellatus*, brain ventricular system at the stage 1-ovl/pec-bud, 165 hpf. (a-d) microCT images, the pseudo-colour volume rendering of the head region overlayed with a surface view of the manually-segmented brain ventricles. (a-d) dorsal, lateral, ventral, and rostral views, respectively. The inset in the upper right corner in (b) illustrates the median ventricular sulcus (MVS) branches off from the ventral bottom of the median ventricle of the mesencephalon. For annotations, see List of Abbreviations. Scale bars = 100 μm.

#### STAGE 1-ovl/pec-bud (165 hpf)

In the Rosy Bitterling, the ventricular expansion is completed at stage 165 hpf (compare Figure 1a, b). The rhombencephalic ventricle (RVe) is diamond-shaped (rhombic) in dorsal and rostral aspects (Figure 3a and c). The bilateral tectal ventricles (TeVe) resemble scallop shells (Figure 3a). The telencephalic ventricle is triangular in rostral aspect (Figure 3d). A pair of oval fossae appear caudal-ventral to the midbrain (Figure 3c, of). These fossae are occupied by the rostral cerebellar thickenings, which develop into the valvula cerebelli of the adult (Wullimann and Knipp, 2000). The lateral recess (LR) and the posterior recess (PR) of the hypothalamic ventricle are in shallow groves (Figure 2a).

#### STAGE pec-bud (185 hpf)

The width of the rhombencephalic ventricle (RVe) decreases to less than half that of the tectal ventricles (TeVe) (Figure 4a). The RVe gradually flattens along its dorsal-ventral axis (compare Figures 3b and 4b). The dorsal surface of the TeVe appears smoother due to the further development of the optic tectum (mammalian superior colliculus) and torus semicircularis (mammalian inferior colliculus; Figure 4a). There is a deep midline ridge separating the left and right parts of the TeVe (Figure 4a, d). The preoptic recess (PoR) is compressed, and reduced in width (compare Figures 3c and 4c). The oval fossae enlarge with the growth of the rostral cerebellar thickening (compare Figures 3c and 4c). The lateral recess and posterior recess become visible and extend outward (Figure 2b).

**Figure 4.**
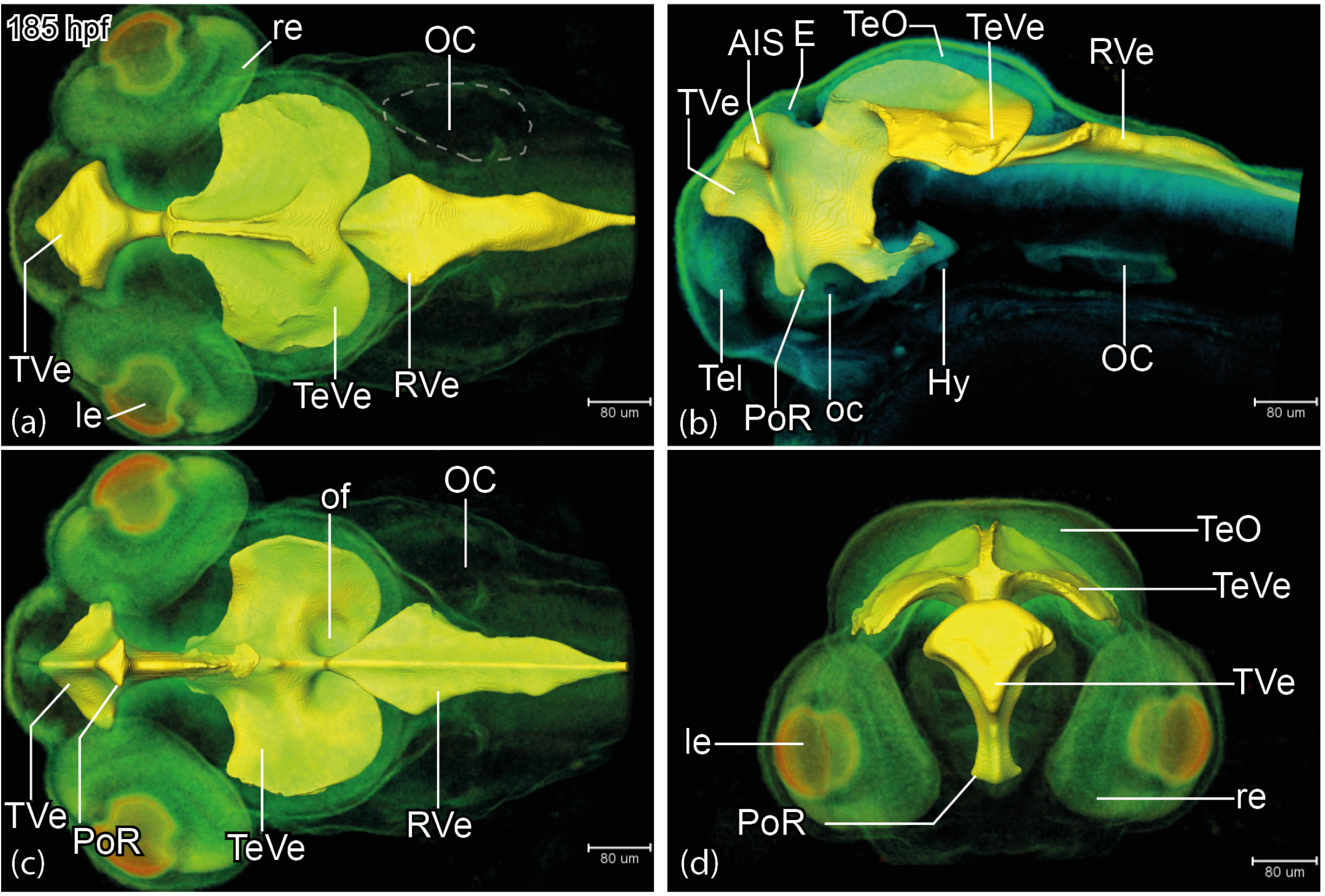
*Rhodeus ocellatus*, brain ventricular system at the stage pec-bud, 185 hpf. (a-d) microCT images, the pseudo-colour volume rendering of the head region overlayed with a surface view of the manually-segmented brain ventricles. (a-d) dorsal, lateral, ventral, and rostral views, respectively. For annotations, see List of Abbreviations. Scale bars = 80 μm.

#### STAGE high-pec (210 hpf)

Compared to earlier stages, the brain ventricles appear compressed (compare Figures 4a and 5a). However, the dorsal ventricle of the anterior intraencephalic sulcus (AIS) appears is expanded posteriorly, probably due to the eversion of the telencephalic ventricle (TVe). The tectal lobes grow larger and appear to fuse in the midline; as a result, the deep midline ridge of the TeVe appears compressed rostrally (Figure 5b). The lateral recess (LR) extends basalward to form a flange at the posterior recess (PR). The inferior lobe forms around the lateral recess (IL, Figure 2c; Bloch et al. 2019).

**Figure 5.**
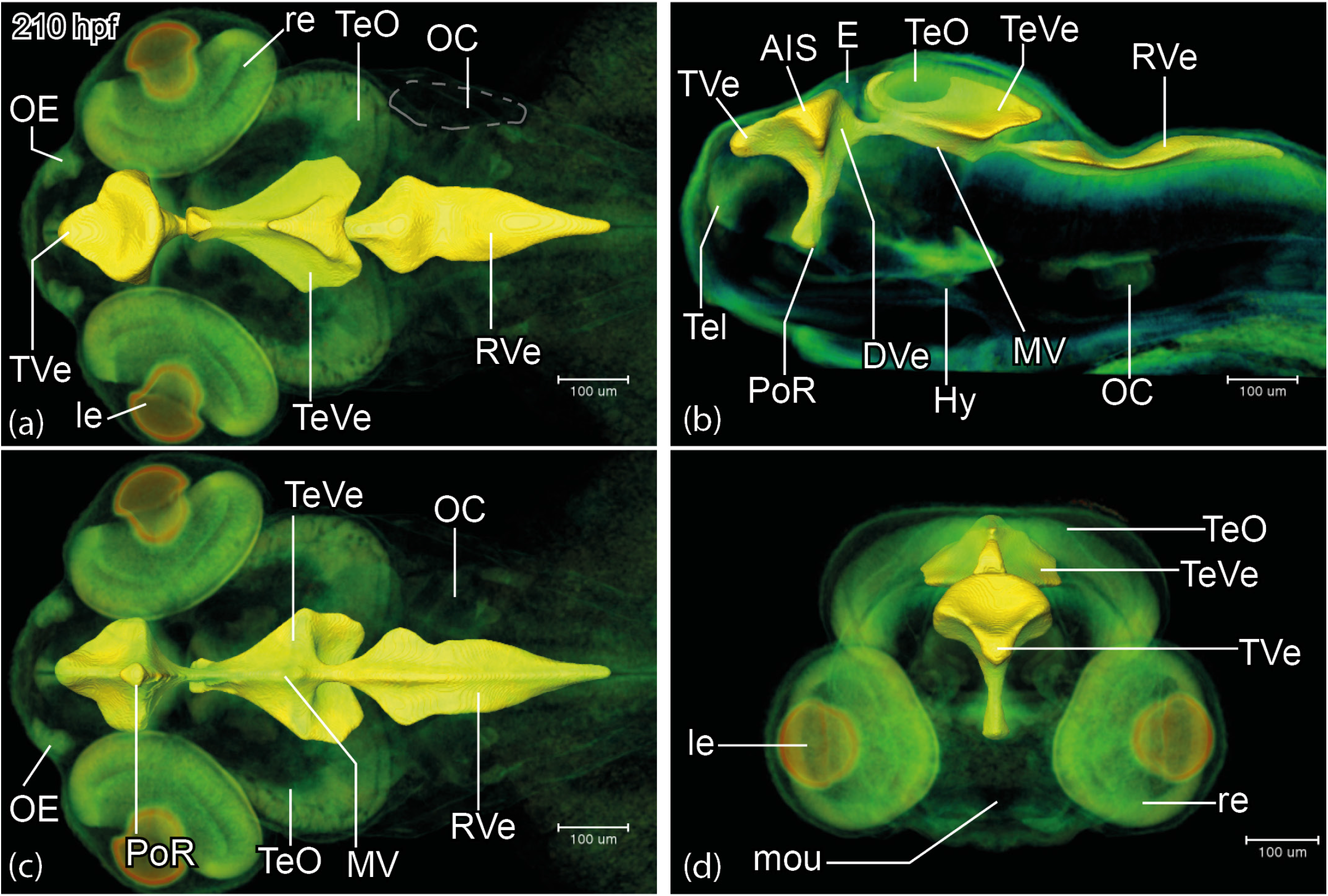
*Rhodeus ocellatus*, brain ventricular system at the stage high-pec, 210 hpf. (a-d) microCT images, the pseudo-colour volume rendering of the head region is overlayed with a surface view of the manually-segmented brain ventricles. (a-d) dorsal, lateral, ventral, and rostral views, respectively. For annotations, see List of Abbreviations. Scale bars = 100 μm.

#### STAGE long-pec (235 hpf)

The rhombencephalic ventricle (RVe) can be divided into two parts, a rostral rhomboid opening with very thin roof plate over it, and a caudal, elongated ventricle between the rhombic lips (RL; Figure 6a). The rostral aspect of the telencephalic ventricle (TVe) gradually deepens from a triangle into a T-shape (Figure 6d). Compared to earlier stages, the tectal ventricles (TeVe) are further compressed and separated from the median ventricle (Figure 6b). The lateral recess (LR) extends more basalward (compare Figures 2c and d), and the inferior lobe (IL) is considerably enlarged. The posterior recess (PR) extends alarward (Figure 2d).

**Figure 6.**
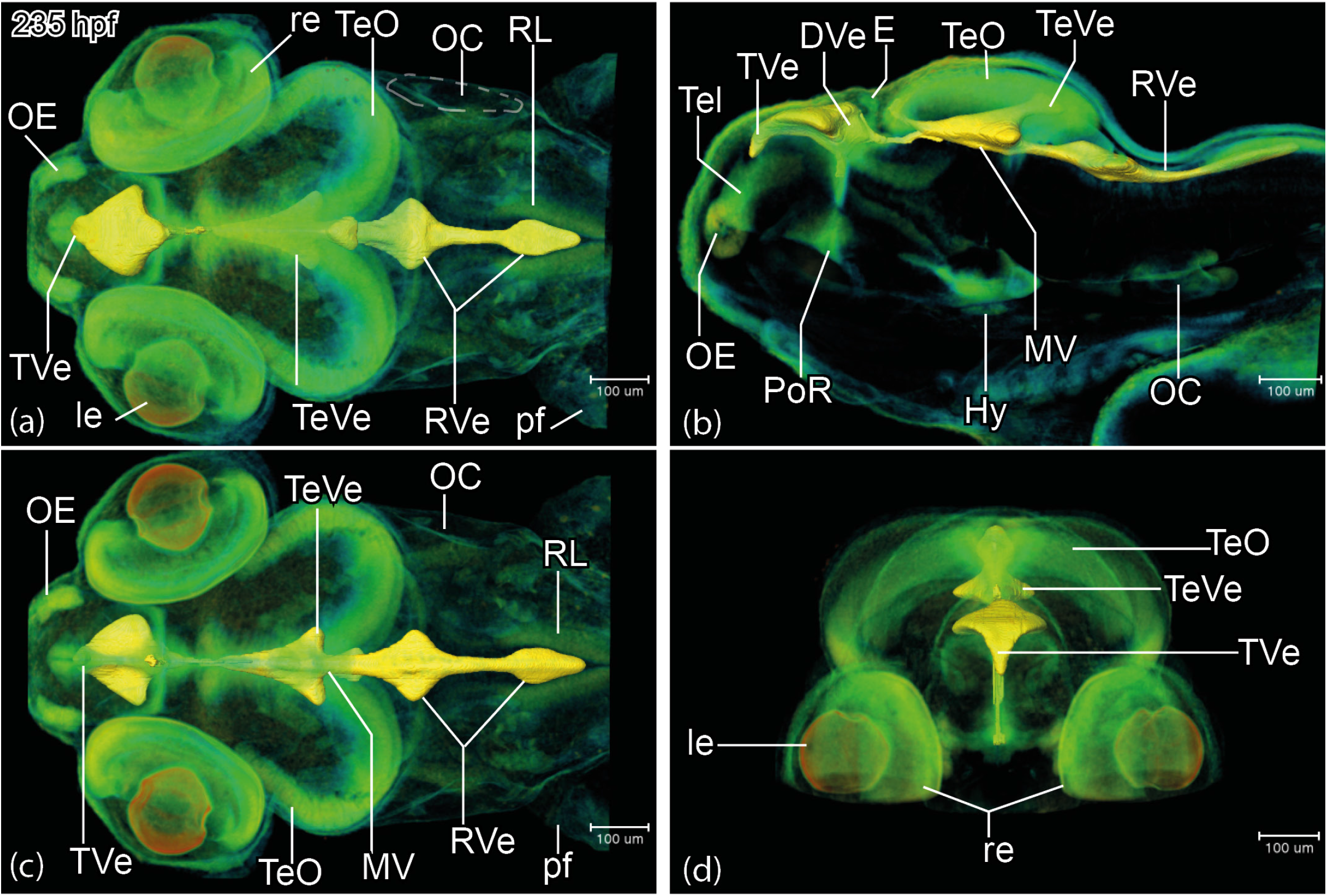
*Rhodeus ocellatus*, brain ventricular system at the stage long-pec, 235 hpf. (a-d) microCT images, the pseudo-colour volume rendering of the head region is overlayed with a surface view of the manually-segmented brain ventricles. (a-d) dorsal, lateral, ventral, and rostral views respectively. For annotations, see List of Abbreviations. Scale bars = 100 μm.

### B. Developmental atlas of the Rosy Bitterling brain

We observed cell clusters in periventricular locations that appeared brighter than the adjacent gray matter (eg. Figure 7j). The distribution of these cell clusters is highly consistent with the distribution of proliferation zones detected during neurogenesis in the zebrafish (Mueller and Wullimann, 2005, 2016). This fact has helped us to map putative proliferation zones of the developing bitterling brain and delineate brain territories. As a consequence, our delineation of anatomical structures is based on three types of observation: (i) their topological relationship to proliferation zones; (ii) their relative location to brain ventricles (see previous section) and other landmarks such as commissures and fiber tracts; (iii) their grayscale values in virtual microCT sections. To take an example, the zona limitans intrathalamica (Zli) can be demarcated by its dark appearance in contrast to the surrounding, bright-white thalamic tissues (Figure 7g, h).

**Figure 7.**
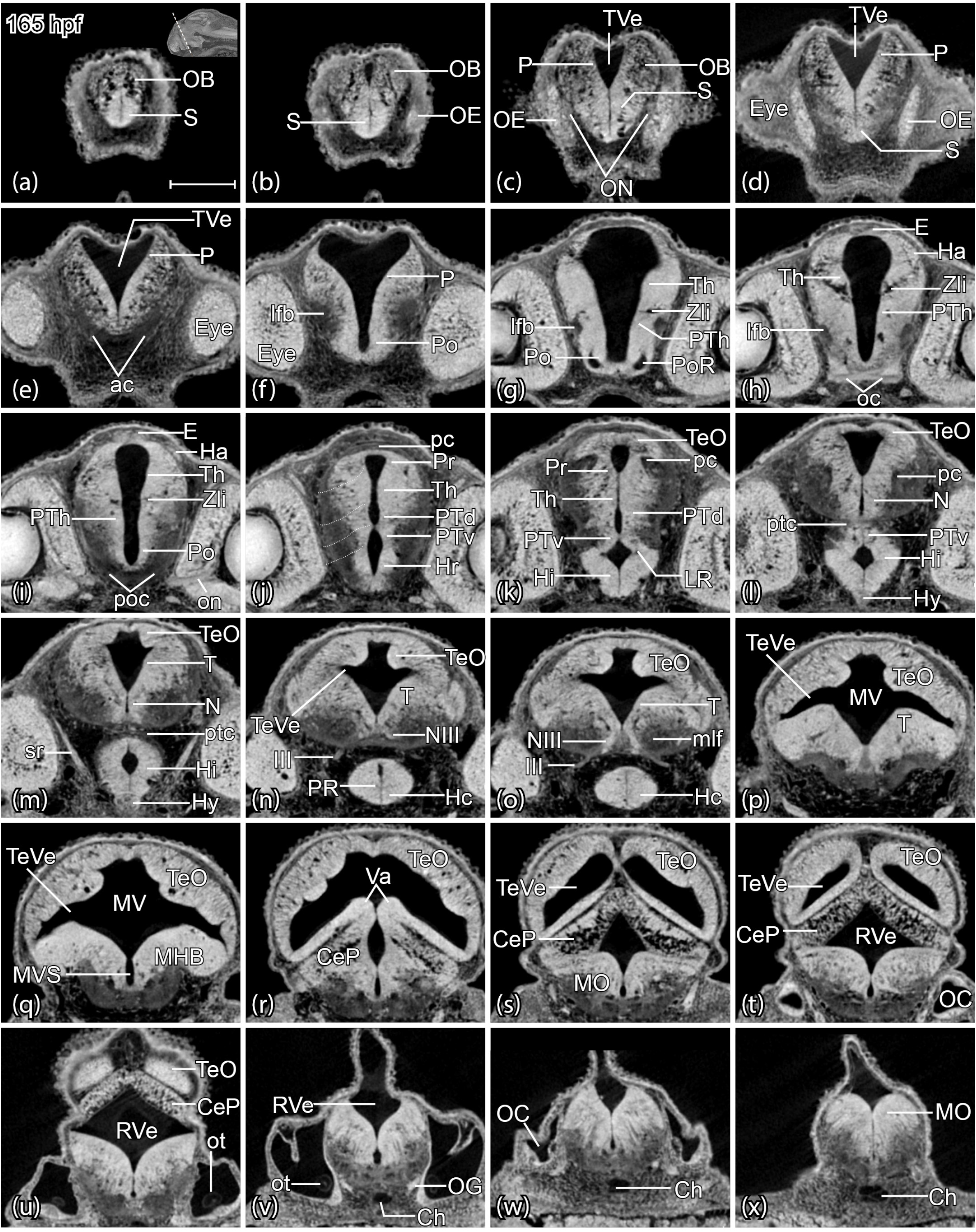
*Rhodeus ocellatus*, brain cross-sectional anatomy, stage 1-ovl/pec-bud, 165 hpf. (a-x) microCT images, virtual sections, transverse plane, dorsal towards the top, sections from rostral to caudal, direction of section plane indicated in inset in (a). For annotations, see List of Abbreviations Scale bars = 100 μm

#### Secondary prosencephalon

Rostral to the telencephalic region, the olfactory epithelium (OE) is visible as a very bright region from the 1-ovl/pec fin stage (165 hpf, Figure 7a). At the long-pec stage (235 hpf) the olfactory epithelium develops into a bow-shaped structure surrounding the lumina of the olfactory pits (Figure 10a). The olfactory epithelium (OE) is connected to the olfactory bulb (OB) through the easily-recognizable olfactory nerve (ON; Figures 7c, 9b and 10b). The olfactory bulb (OB) is characterized by its glomerular structure (Figure 7a, 8a, 9a and 10a; Dynes and Ngai 1998).

**Figure 8.**
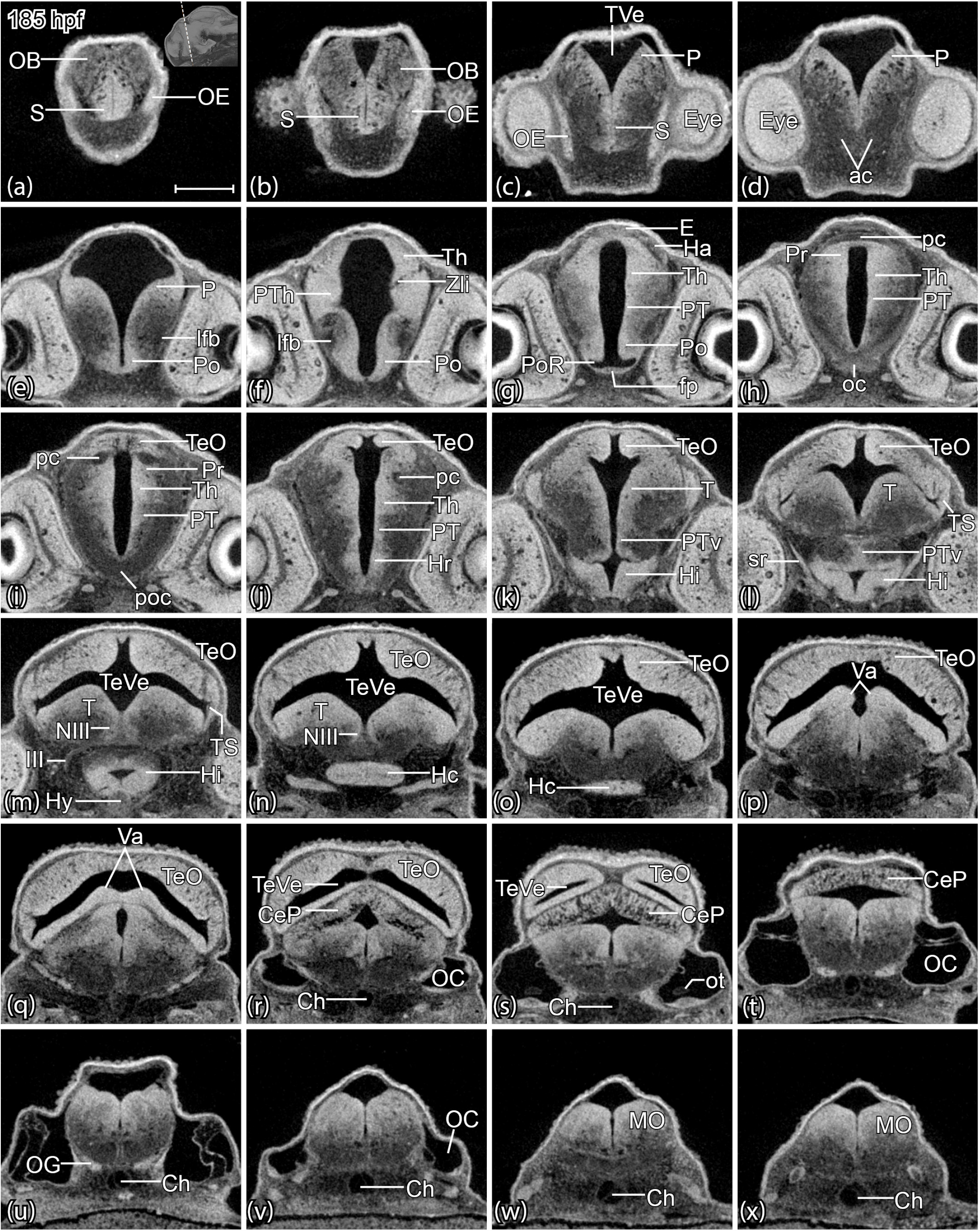
*Rhodeus ocellatus*, brain cross-sectional anatomy, stage pec-bud, 185 hpf. (a-x) microCT images, virtual sections, transverse plane, dorsal towards the top, sections from rostral to caudal, direction of section plane indicated in inset in (a). For annotations, see List of Abbreviations. Scale bars = 100 μm.

**Figure 9.**
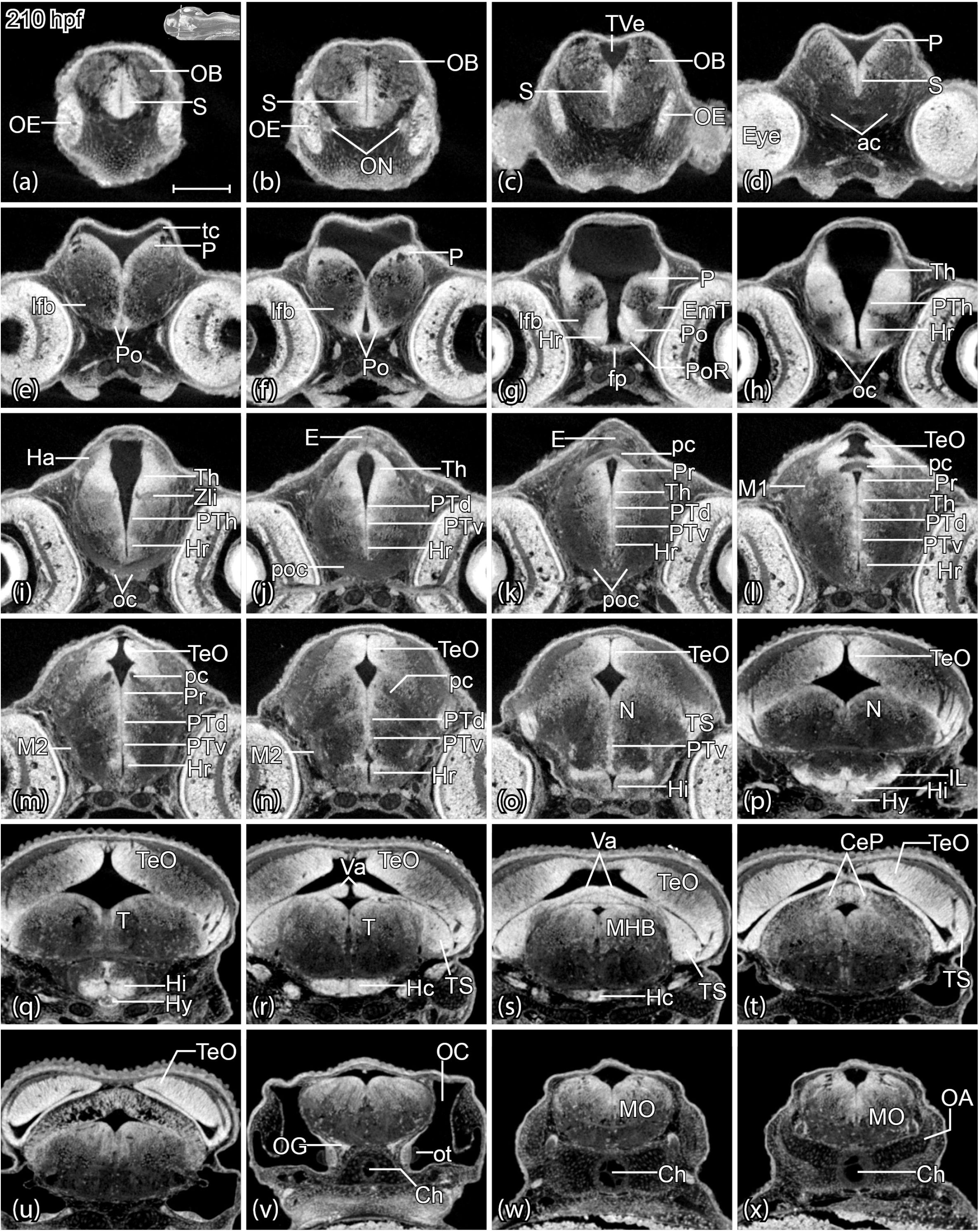
*Rhodeus ocellatus*, brain cross-sectional anatomy, stage high-pec, 210 hpf. (a-x) microCT images, virtual sections, transverse plane, dorsal towards the top, sections go from rostral to caudal, direction of section plane indicated in inset in (a). For annotations, see List of Abbreviations. Scale bars = 100 μm.

**Figure 10.**
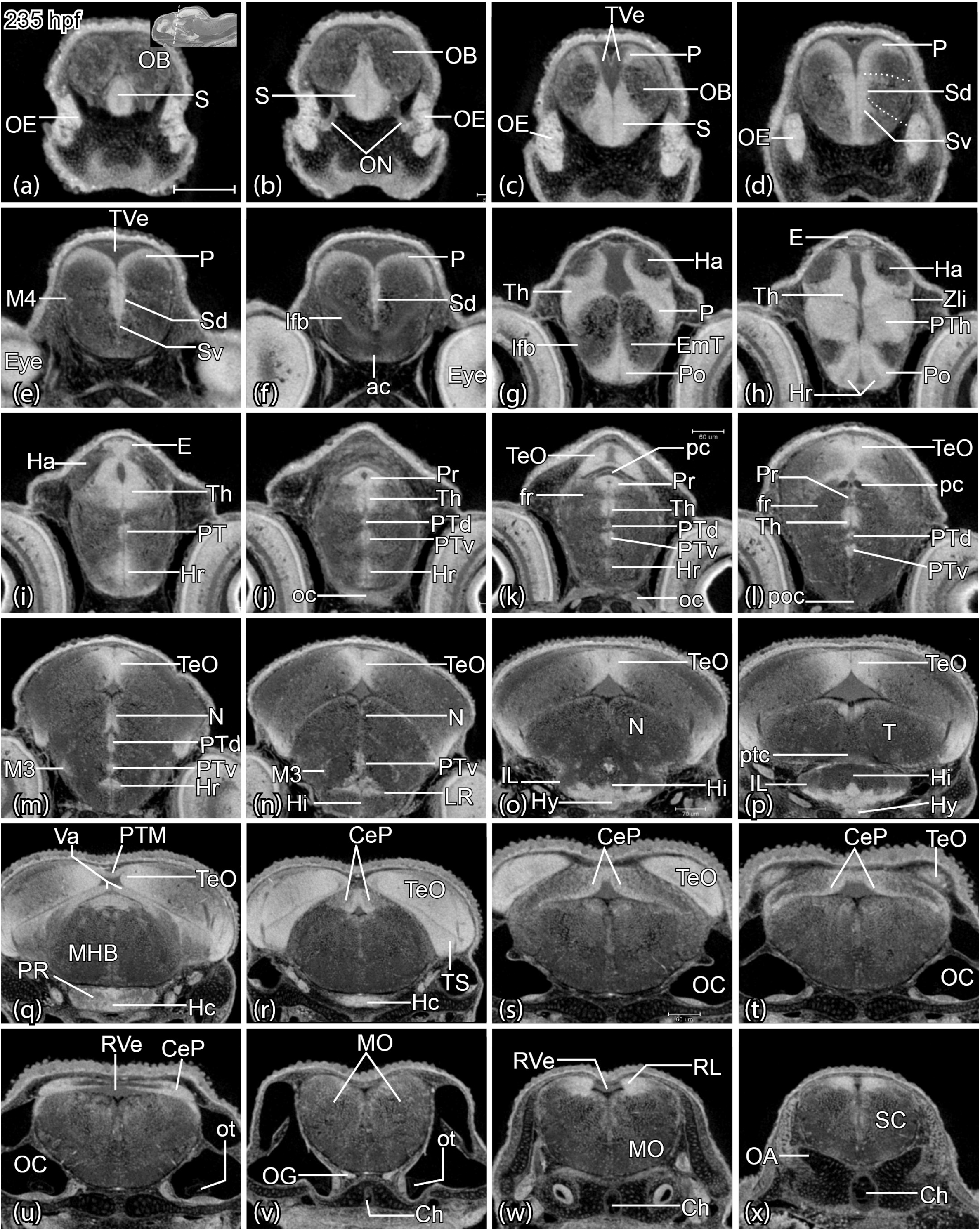
*Rhodeus ocellatus*, brain cross-sectional anatomy, stage long-pec, 235 hpf. (a-x) microCT images, virtual sections, transverse plane, dorsal towards the top, sections go from rostral to caudal, direction of section plane indicated in inset in (a). For annotations, see List of Abbreviations. Scale bars = 100 μm.

The telencephalic ventricle is located at the dorsum of the pallium and the tela choroidea is visible in its roof. The periventricular proliferation zones of the subpallium (S) and pallium (P) appear bright in virtual transverse sections. At the long-pec stage (235 hpf), the bright subpallial cell clusters were separated by distinct, dark boundaries (dashed line in Figure 10d), corresponding to the dorsal and ventral subdivisions of the subpallium (Sd and Sv). The zones of pallial proliferation consist of a few cell rows, and are visible as a pair of arches flanking the dorsal subpallium (Figure 7c, 8c, 9d, 10c). The telencephalic migrated area (M4) was recognizable in the form of cell clusters at the margin of the lateral subpallium (Figure 10e).

Immediately caudal to the subpallium, the anterior commissure (ac) and lateral forebrain bundle (lfb) appeared as dark fiber bundles crossing the rostral end of the forebrain (Figure 7e,f, 8d,e, 9d,e, 10f). We shall define the anterior commissure (ac) and the postoptic commissure (poc) as the boundaries of the preoptic region (Po). This region recently has also been designated as the alar hypothalamus (aHyp) or the optic recess region (ORR) (Affaticati et al., 2015; Schredelseker and Driever, 2020; Slack, 2005). The proliferation zone of the preoptic region (Po) was easy to identify as a triangular region surrounding the optic recess. The Po proliferation zone is broad on its ventral side and thins out dorsally (Figure 7f, 8e, 9e, 10g). The optic chiasma (oc) which has a bright, thick appearance, decussates in the midline (Figure 7h, 8h, 9h, 10j); it marks the anterior end of the neural axis.

In the basal hypothalamic region, the hypophysis is located topologically acroterminal (anterior) to the hypothalamus (Figure 2). It projects form the ventral midline of the brain (Figure 2). As in the zebrafish developmental brain atlas (reference?), we divide the basal hypothalamus into the following regions (i) the rostral hypothalamus (Hr); (ii) the caudal hypothalamus (Hc); and between them: (iii) the intermediate hypothalamus (Hi), located near the hypophysis, and including the inferior lobe (IL). The hypothalamus encloses the hypothalamic ventricles, including the lateral recess (LR) in the Hi, and the posterior recess (PR) in the Hc. This division is consistent with those classically used for describing the zebrafish hypothalamus (Wullimann et al., 1996; Manoli and Driever, 2014; Biran et al., 2015; Mueller and Wullimann, 2016; Muthu et al., 2016).

Recent molecular studies in the zebrafish have improved the interpretation of the teleostean hypothalamus and its evolutionary relationships with the mammalian hypothalamus (Baeuml et al., 2019; Herget et al., 2014). Likewise, gene expression in the PR tuberal region of the embryonic zebrafish has revealed homology with two domains of the mammalian hypothalamus: the TuV (tuberal region, ventral part) and TuI (tuberal region, pars intermedia). Therefore, Schredelseker and Driever (2020) proposed to refer to this region as PRR (posterior recess region) to set it apart as a teleost-specific entity. Because our analysis of the Rosy Bitterling is based on anatomical microCT data, are not able to confidently relate our findings those just cited. We therefore look forward to molecular studies in bitterlings.

#### Diencephalon

The prosomeric model divides the diencephalon into alar plate and basal plate derivatives. In this model, the pretectum (aP1), the thalamus proper (aP2), and the prethalamus (aP3) form the alar plate portions of the diencephalon (Lauter et al., 2013). In contrast, the proliferation zones of the nucleus of the medial longitudinal fasciculus (N; bP1), and the dorsal (bP2) and ventral (bP3) posterior tegmentum (PT; Mueller and Wullimann, 2005, 2016) form the corresponding basal plate derivatives.

The most prominent structure in the diencephalic roof plate is the epiphysis (E), a swelling in the midline (Figure 7h, 8g, 9j, 10i). The habenular nuclei (Ha) are located one on each side of it (Figure 7h, 8g, 9i, 10i) and show discrete habenular cell clusters from the pec-bud stage (185 hpf) onwards (Figure 8g). Notice that the fasciculus retroflexus (fr) appeared in our microCT photographs in the form of distinctive, dark fiber-bundles in the gray matter (Figure 10k, i). The fasciculus originates from the habenular nuclei and connects to the interpeduncular nucleus across the isthmus (r0) and rhombomere 1 (Akle et al., 2012). In the prosomeric model, the fasciculus retroflexus is used to delimit pretectum (P1) and thalamus (P2; Akle et al., 2012; Lauter et al., 2013; Puelles, 2019). We use the caudal end of the posterior commissure as the caudal boundary of the pretectum (P1), which divides the diencephalic and mesencephalic areas (Figures 7j, 8h, 9l, 10k).

In the thalamic region, we identified the zona limitans intrathalamica (Zli) as a dark band (Figure 7i, 8f, 9i, 10h) that marks out the boundary between the prethalamus (P3) and thalamus (P2). Therefore, we annotated the separate periventricular proliferation zones of the prethalamus (PTh) and thalamus (Th) based on their topological relationships (anterior vs. posterior) and Zli landmark (e.g. Figure 10j).

The thalamic eminence (EmT) is a relatively complex region of the diencephalon and is viewed in the prosomeric model as the anterior portion of the prethalamus (and is therefore often termed the “prethalamic eminence”, PThE). However, while the EmT (or PThE) generates glutamatergic derivatives, the prethalamus proper mostly contains GABAergic neurons. In addition, some recent studies in the zebrafish and tetrapods indicated that the Emt/PThE contributes to telencephalic territories such as the medial extended amygdala and the newly-identified nucleus of the lateral olfactory tract (Alonso et al., 2020; Porter and Mueller, 2020; Vicario et al., 2017). Due to the lack of gene expression data in bitterlings, we are conservative in our analyses and place the EmT topologically anterior to the prethalamus, and posterior to the preoptic region, similar to the condition described for larval zebrafish (Wullimann and Mueller, 2004). It abuts the lateral forebrain bundle (Mueller, 2012) which is identifiable in our CT scans (Figure 7f, 8e, 9e, 10f). The use of molecular markers such as Tbr-1 (Wullimann, 2009; Wullimann and Mueller, 2004) would help validate the anatomical hypotheses in the present study.

We conclude this section by noting that the PT has distinct PTd and PTv proliferation zones (eg. Figure 10k). Because of the cephalic flexure, the dorsal-ventral topology in the virtual transverse sections, corresponds to the anterior-posterior axis of the neural tube.

#### Mesencephalon and rhombencephalon

The boundary between the diencephalon and mesencephalon is defined dorsally by the posterior end of the posterior commissure, and ventrally by the anterior margin of the oculomotor nerve root (Moreno et al., 2016). In the tegmentum, we were able to identify the oculomotor nerve (III), which projects ventrolaterally from the oculomotor nucleus (NIII) and exits from the ventral surface of the brain (e.g., Figure 7n and o). The course of the oculomotor nerve is a useful landmark for helping to identify the oculomotor nucleus (NIII) and the basal plate of the mesencephalon.

In the prosomeric model, the mesencephalon contains two mesomeres, m1 and m2 from anterior to posterior. The tectal gray, optic tectum (mammalian superior colliculus), and torus semicircularis (mammalian inferior colliculus) constitute the alar plate of m1. The oculomotor nucleus (NIII) represents the basal plate of m1.

We noticed two pairs of tectal membrane thickenings which invaginate into the tectal ventricle, toward the tegmentum, at the 1-ovl/pec-bud stage (165 hpf; Figures 7q and r). During development, the boundaries between these thickenings gradually disappears as they growth together (Figure 8o and 9o). The tectal proliferation zones are distinct. At the 1-ovl/pec-bud stage (165 hpf) and the pec-bud stage (185 hpf), the tectal region appears as a large, bright field with microCT (Figure 7o and 8l). Beginning at the high-pec stage (210 hpf), the rostral tectal proliferation becomes restricted to one mediodorsal cluster and two bilateral clusters (Figure 9o and 10n). These lateral and medial proliferation zones merge in the midline at caudal levels and form a continuous cap of tectal proliferation (e.g., Figure 9t).

The torus longitudinalis (TL) is a specialized brain region exclusive to ray-finned fish (Folgueira et al., 2020; Wullimann, 1994). We identified the TL from rostral to caudal along the medial margins of the optic tectum. Virtual horizontal sections through the optic tectum horizontally from the level of the epiphysis and the rostral cerebellar thickening, showed the TL at the top of the mesencephalic ventricle (Figure 11).

**Figure 11.**
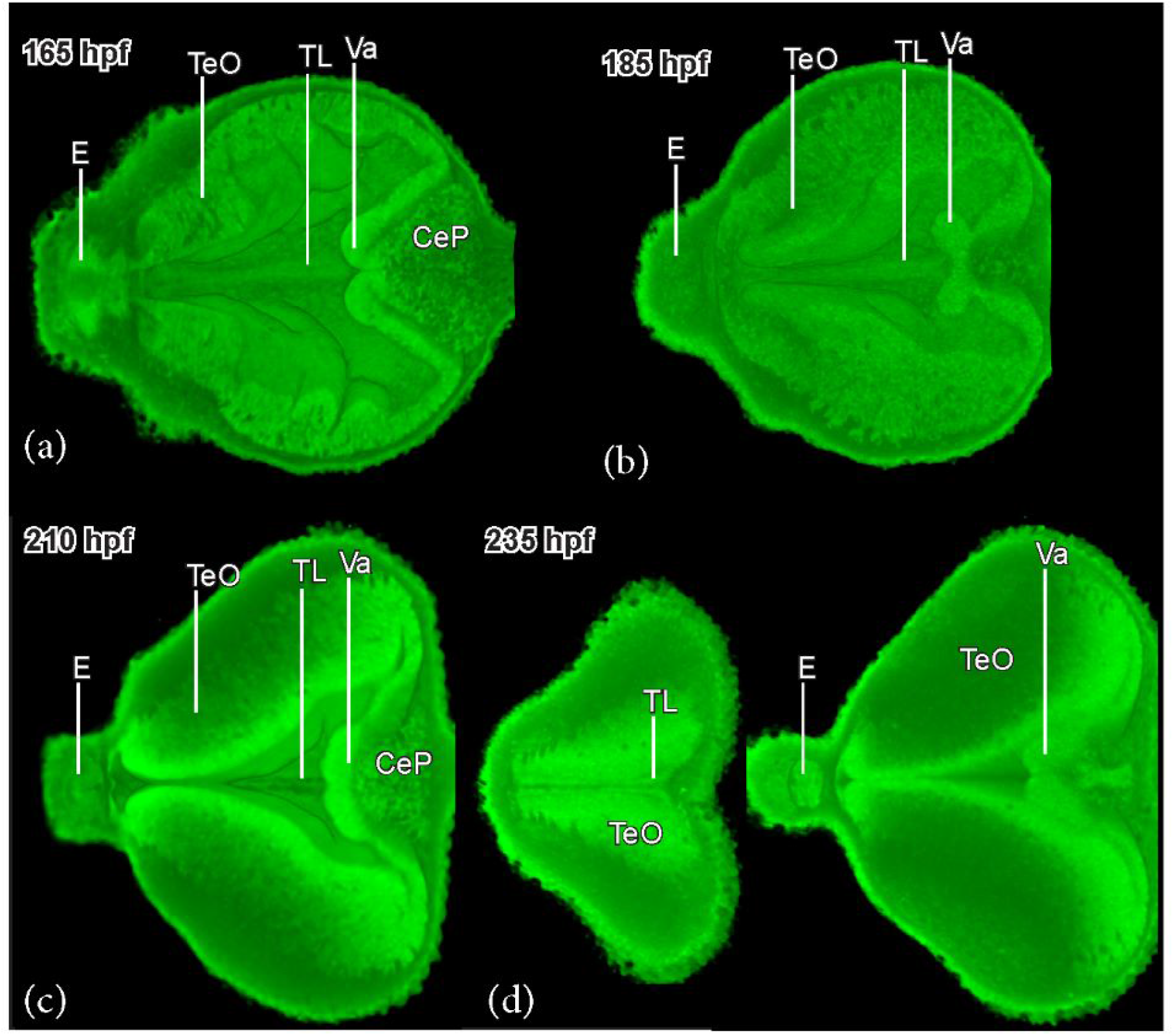
*Rhodeus ocellatus*, development of the torus longitudinalis. Virtual dissection, head to the left. (a) stage 1-ovl/pec-bud, 165 hpf. (b) stage pec-bud, 185 hpf. (c) stage high-pec, 210 hpf. (d) stage long-pec, 235 hpf. For annotations, see List of Abbreviations.

The r0, or isthmus, is coincident with the midbrain-hindbrain boundary (MHB). The MHB is composed of the posterior tectal membrane and the rostral cerebellar thickenings (the valvula cerebelli; Wullimann and Knipp, 2000). The rostral cerebellar thickenings appeared bright throughout the developmental period in this study (Figure 7r, 8q, 9r, 10q). The cerebellar plate appeared bright only at its basal and medial aspects (e.g., Figure 10t and u). The trochlear nucleus (NIV) topologically belongs to r0. It is easier to identify the trochlear decussation (DIV) and the commissura cerebelli (Ccer) in the midsagittal plane (Figure 12a-d) in the valvula cerebelli (Va) than in the horizontal or the transverse planes. Then follow the caudolateral projection of the trochlear axon in the horizontal plane (Figure 12e) until it exits the brain as the trochlear nerve (IV, Figure 12f) between the torus semicircularis and rhombencephalon. The axonal tract of the trochlear nucleus delineates the boundary between r0 and r1.

**Figure 12.**
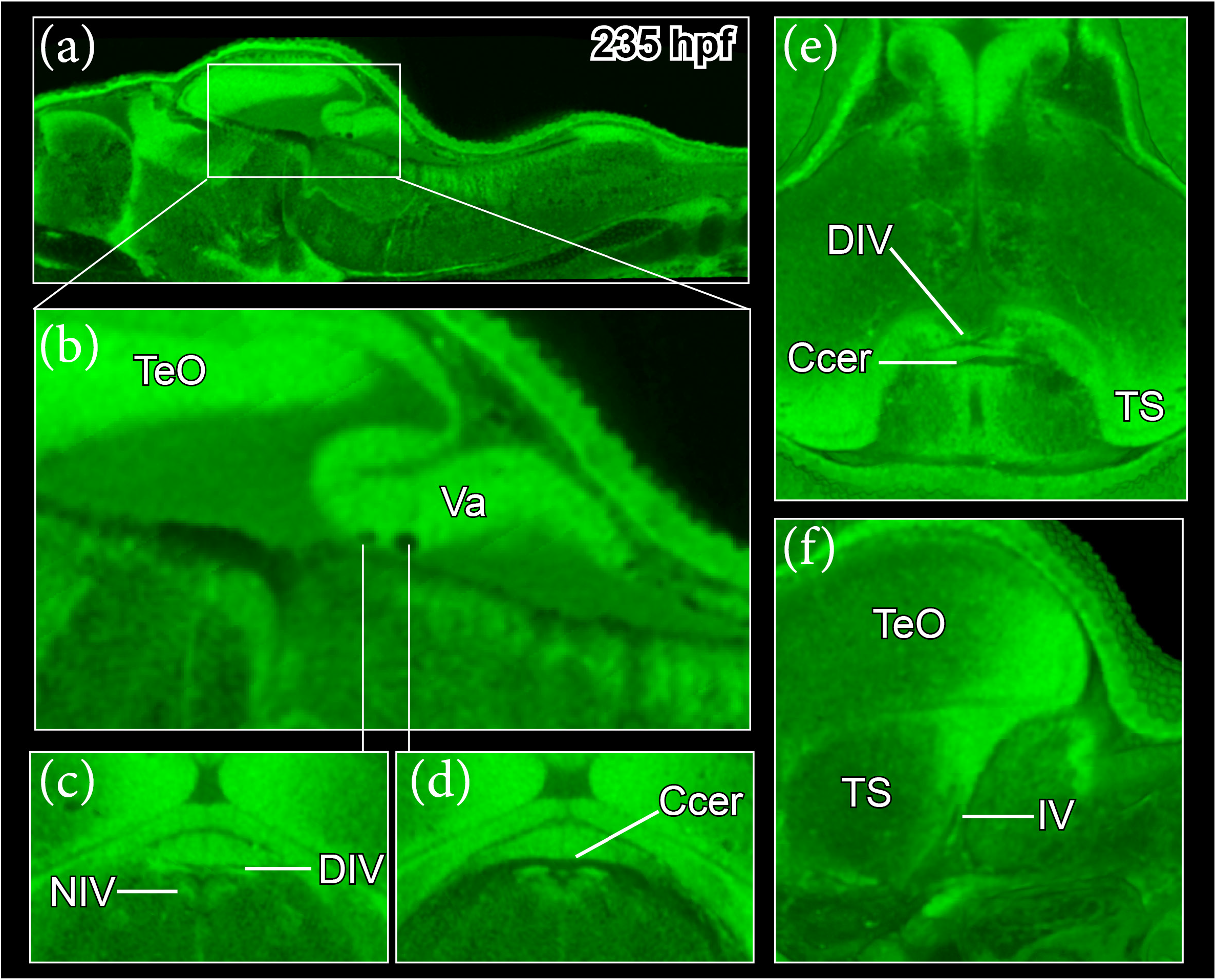
*Rhodeus ocellatus*, trochlear nerve, stage long-pec, 235 hpf. (a-f) microCT images, virtual sections. (a and b) midsagittal plane, head to the left, dorsal towards the top. (c and d) transverse plane, dorsal towards the top. (e) horizontal plane, head towards the top. (f) parasagittal plane, head to the left, dorsal towards the top. For annotations, see List of Abbreviations.

The fasciculus retroflexus (fr) of teleosts projected to the interpeduncular nucleus (NIn) and so we used it to identify the NIn in the bitterling embryo (Figure 13a). According to Lorente-Cánovas et al. (2012), the interpeduncular nucleus is at the basal plate at the level of the isthmus (r0) and r1. Rhombomere r1 is devoid of cranial motor neurons (Nieuwenhuys and Puelles, 2016), so we used the posterior margin of the interpeduncular nucleus as a landmark for the boundary between r1 and r2.

**Figure 13.**
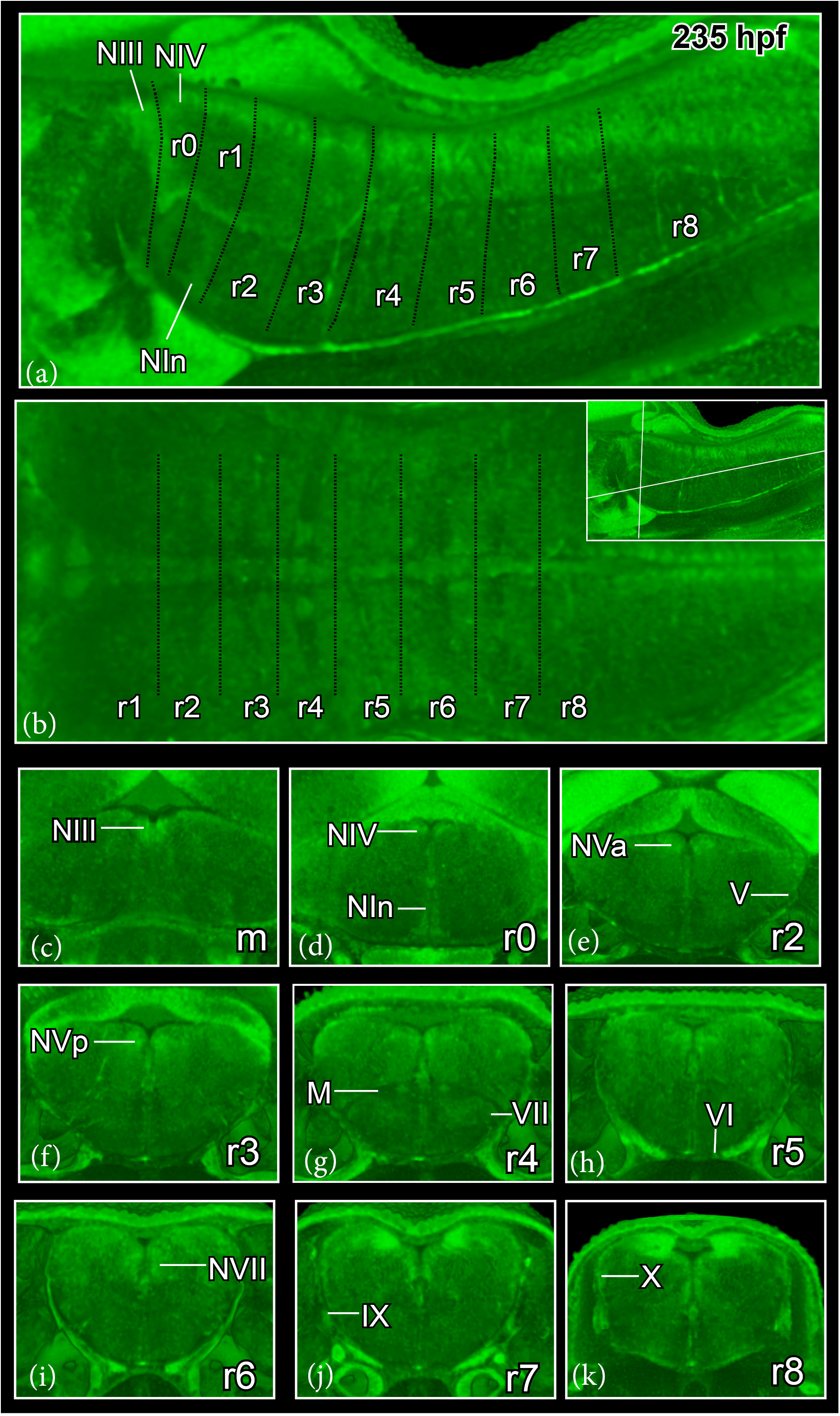
*Rhodeus ocellatus*, hindbrain segmentation, stage long-pec, 235 hpf. (a) microCT images, virtual sections, midsagittal plane, dorsal towards the top, head to the left. The black dash line indicates the rhombomeric boundaries. (b) horizontal plane, head to the left, direction of section plane indicated in inset at the upper right corner. (c-k) transverse plane, dorsal towards the top, sections go from the rostral to caudal, direction of section plane indicated in inset in (b). For annotations, see List of Abbreviations.

In the more caudal rhombencephalic region, the roof plate is a thin layer of tela choroidea, which is visible in microCT scans because it remains intact during the fixation and staining procedures (e.g., Figure 7x). The dorsal medullary proliferation zone is broad, and expands ventrally up to the high-pec stage (210 hpf; (Figure 7v, 8v, and 9v). At the long-pec stage (235 hpf) it becomes more spatially restricted, forming the rhombic lip proliferation zone (Figure 10w).

The boundaries between rhombomeres from r2 to r8 are visible in early bitterling embryos at 135 hpf (Figure 14), but soon become less visible from the 1-ovl/pec fin stage (165 hpf). Teleosts retain a segmented pattern of reticulospinal neurons through embryonic life to adulthood (Gilland et al., 2014). For example, the large bilateral Mauthner neurons (M) are markers of r4 (Eaton and Farley, 1973; Moens and Prince, 2002). By virtually sectioning the 135 hpf embryo at the axial level of r4, we determine that the Mauthner cell reside near the center of rhombomere 4, rather than at the segmental boundaries (Figure 14). The rhombomeric segments of the older embryo (235 hpf, Figure 13a to k) were thereby identified, by assuming that each reticulospinal neuronal cluster is located in the center of each rhombomeres.

**Figure 14.**
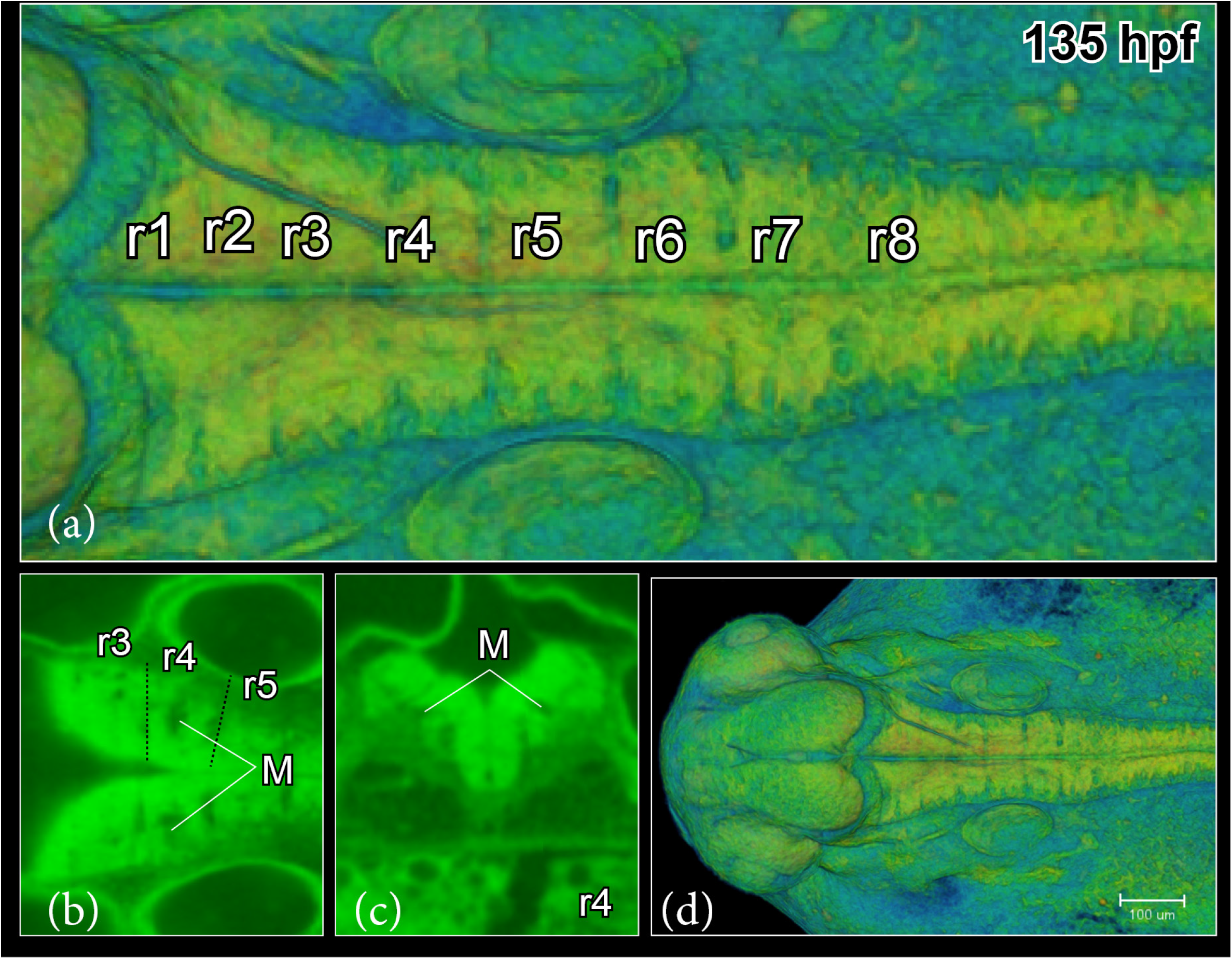
*Rhodeus ocellatus*, hindbrain segmentation, 135 hpf. (a and d) microCT images, volume rendering, show boundaries between rhombomeres. (b) virtual section, horizontal plane, head to the left, show the location of the Mauthner cell at the centre of r4. (c) virtual section, transverse plane, dorsal towards the top. For annotations, see List of Abbreviations.

The nerve roots of the cranial nerves are also reliable landmarks of rhombomeres. By tracing the projection of cranial nerves, we identified the trigeminal nerve root (V) in r2 (Figure 13e), the facial root (VII) and the accompanying vestibulo-cochlear nerves associated with r4 (Figure 13g), the abducens root (VI) in r5 (Figure 13h), the glossopharyngeal root (IX) in r7 (Figure 13j), and the vagus root (X) in r8 (Figure 13k). Rhombomere r2 was also indicated by cell bodies of the anterior trigeminal motor neurons (NVa, Figure 13e) and r3 by the posterior trigeminal motor neurons (NVp, Figure 13f). Facial nerve motor neurons (NVII) are distinct in r6 (Figure 13i).

## DISCUSSION

### Brain imaging and the 3D visualization of neuroanatomy

To gain insight into the complex morphogenesis of the Rosy Bitterling brain, we analyzed the formation of the brain ventricular system in 3D from the 1-ovl (150 hpf) stage to the long-pec (235 hpf) stage. We combined annotations of brain functional subdivisions and morphological landmarks, based on microCT scanning results and on the literature for the zebrafish embryo. We found that 3D visualization with microCT scanning protocols were extremely useful for the study of the bitterling brain. They provide an updating of imaging modalities for morphological and anatomical analyses in this non-model organism. This systematic application of microCT may help provide an essential foundation for future comparative studies of the teleost brain.

Our study provides 3D reconstructions of the brain ventricles in the bitterling; they show a similar organization to the zebrafish larval ventricular system, as visualized by dye-injection into the hindbrain ventricle (Lowery and Sive, 2005). We have also defined boundaries of brain ventricular subdivisions, based on published anatomical studies (Turner et al., 2012). Furthermore, we found that the flexure of the neuraxis is easily visualized continuously from the rostral tip of the brain to the spinal cord. Understanding the cephalic flexure is crucial, due to its profound effect on the topology of the brain, especially in the highly complex secondary prosencephalon (Hauptmann and Gerster, 2000; Puelles, 2019). It has been shown that the mechanism of brain ventricle development is highly conversed among vertebrates (Lowery and Sive, 2009). The cerebrospinal fluid in the brain ventricular system could contribute to specialization of the early brain, because the production and circulation of cerebrospinal fluid performs an important role in the homeostasis of the central nervous system (Fame et al., 2016).

In contrast to other vertebrates, teleosts and other actinopterygian fish, lack a pair of lateral ventricles in the telencephalon (Wullimann and Rink, 2002). Instead, they show a T-shaped midline telencephalic ventricle that is the result of a morphogenetic process called ‘eversion’ (Mueller and Wullimann, 2009). Our 3D models of the Rosy Bitterling brain show that the anterior intraencephalic sulcus (AIS) develops much earlier than the eversion of the telencephalic ventricle, whereas the expansion of the dorsal ventricular surface of the AIS is synchronous with the eversion process. Our results are consistent with the pattern of telencephalic ventricle morphogenesis described in the zebrafish (Folgueira et al., 2012). We were able to visualize the development of the deep ventricular sulcus (AIS), followed by an anterolateral eversion of the telencephalic neuroepithelium.

We found distinct periventricular cell clusters in the gray matter, with higher greyscale values than the surrounding tissue. Their distribution pattern was consistent with the distribution of proliferation zones described during neurogenesis of zebrafish (Mueller and Wullimann, 2003; Mueller and Wullimann, 2016; Mueller et al., 2006; Wullimann, 2009; Wullimann and Knipp, 2000; Wullimann and Mueller, 2004). It is possible that newly postmitotic neurons appear brighter than most of the postmitotic cell masses in more peripheral positions, remote from the periventricular proliferation zones. The results of immunohistochemistry in the zebrafish for the neurotransmitter GABA (γ-aminobutyric acid) in zebrafish (Mueller and Wullimann, 2016; Mueller et al., 2006; Panganiban and Rubenstein, 2002), and PCNA (proliferation cell nuclear antigen) for proliferation patterns (Wullimann and Knipp, 2000; Wullimann and Mueller, 2004; Wullimann and Puelles, 1999) are consistent with our interpretation of the proliferation zones.

Our study will hopefully pave the way for more detailed analyses of the bitterling brain. We hope that it will also prove valuable in studies using the growing number of fish models (and even non-fish models). Application of microCT 3D imaging provides a heuristic model of the brain, an extremely complex anatomical region. Importantly, our approach is validated by the fact that the profile of CT values displayed here in the bitterling brain are consistent with genoarchitecture identified in previous neurodevelopmental studies. For example, our annotation of the zona limitans intrathalamica (ZLI) is extremely close to the highly conserved expression pattern of the gene sonic hedgehog (*shh*; Vieira et al., 2005; Mueller et al., 2006; Scholpp et al., 2006) in the zebrafish. In addition, microCT allows for the time-efficient imaging of intact brains, while providing a resolution (micron level) sufficient for displaying critical landmarks, groups of neurons such as proliferation zones versus postmitotic cell masses, and white matter tracts. These histological characteristics are critical for detecting landmarks and visualizing structural features as a means to describe developmental neuroanatomy.

The resolution of microCT is inferior to light microscopy, whether it is paraffin histology, or the light sheet microscopy-based imaging of fluorescence-stained intact brains. It is also lower in resolution than Synchrotron imaging, whose resolution reaches the sub-micron level, and therefore permits quantitative histological phenotyping (Ding et al., 2019). Neurons and neuropils in the central nervous system can be selectively stained with the salts of metallic elements including gold, silver, platinum and mercury (Keklikoglou et al., 2019; Mizutani and Suzuki, 2012), which can compensate for the inability to use antibodies in combination with microCT. In summary, MicroCT imaging produces lower resolution than some imaging modalities, but has the special advantage that complex 3D models can be produced without the need for time-consuming reconstruction from histological material. The specimens imaged with microCT can be subsequently run through paraffin histology if needed.

### Comparison of developmental stages between bitterling and zebrafish

The phylogenetic relationship of the Rosy Bitterling and the zebrafish is a very close one (they are both Cypriniformes: Cyprinidae; Mayden et al., 2009). This closeness allows for a relatively straight forward interspecies comparison of their brain development. The process of brain ventricle inflation, flexion of the neuroaxis, and establishment of prosomeric units, happens in bitterling in much the same way as in the zebrafish. Specifically, the 165 hpf bitterling brain and 30 hpf zebrafish brain are in the same process of brain ventricle inflation and have a similar degree of flexion of the neuroaxis. We also found that the stratification of the proliferation zones of the diencephalic region (Pr, Th, PTd, PTv) is similar in the 185 hpf bitterling brain and 36 hpf zebrafish brain.

We have detected distinct developmental heterochronies between bitterling and zebrafish brains. The bitterling shows precocious development of the inferior lobe and basalward extension of the lateral recess in the hypothalamic region (Figure 2): the inferior lobe is visible as early as the long-pec stage (235 hpf in the bitterling (Figure 15); but not until 48 hpf in the zebrafish. The long-pec stage is defined by the development of the pectoral fin bud, when the pectoral fin bud in both species grows significantly, and its height grows to twice the the width at its base (Kimmel et al., 1995; Yi et al., 2021). In the zebrafish, the inferior lobe develops relatively late, at around five days post fertilization (Bloch et al., 2019). In teleost fish, the inferior lobe is known as a multisensory integration center that is involved in gustatory (Rink and Wullimann, 1998; Wullimann, 2020), visual (Butler et al., 1991), and octavolateralis system (Yang et al., 2007). Further studies can use the detailed descriptions in our bitterling brain developmental atlas to uncover the underlying regulatory mechanisms that control brain development.

**Figure 15.**
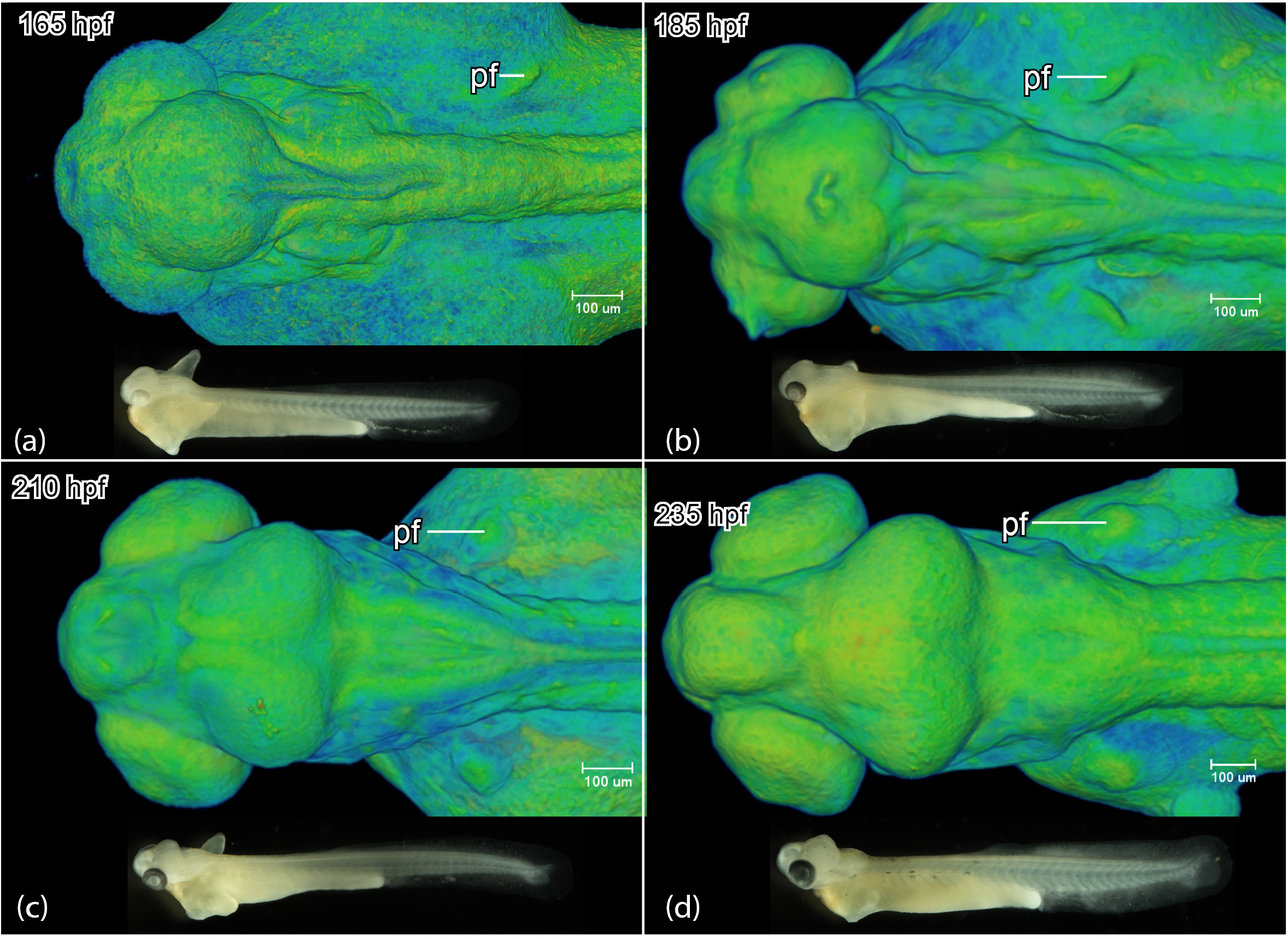
*Rhodeus ocellatus*, development of the brain region and the pectoral fin bud. Dorsal view of the head region microCT images, pseudo-colored volume-renderings, rostral to the left. Lateral view of the embryo, photomicrographs, rostral to the left. (a) 1-ovl/pec-bud, 165 hpf; (b) pec-bud, 185 hpf; (c) high-pec, 210 hpf; (d) long-pec 235 hpf. For annotations, see List of Abbreviations.

## AUTHOR CONTRIBUTIONS

WY and MKR conceived the study. WY performed all neuroembryology studies, including microCT analysis. MR helped with microCT studies. All authors helped WY with interpretation of the microCT data. WY prepared the manuscript and figures.

MKR and TM helped edit the manuscript.

## CONFLICT OF INTEREST

The authors declare that they have no competing interests.

## ACKNOWLEDGMENTS

We are very grateful to Bertie Joan van Heuven (Naturalis Biodiversity Center) for assistance with the microCT, Merijn de Bakker (Leiden University) for laboratory assistance, and Gerda Lamers (Leiden University) for help with sample staining. We thank Doc. RNDr. Martin Reichard (Institute of Vertebrate Biology, Czech Academy of Sciences) for providing the parental Rosy Bitterling fish and assisting us in obtaining embryos through *in vitro* fertilization technology. Wenjing Yi thanks the China Scholarship Council for support (Award no. 201406760046). Thomas Mueller was supported by the Cognitive and Neurobiological Approaches to Plasticity (CNAP), Center of Biomedical Research Excellence (COBRE) of the National Institutes of Health (NIH grant number P20GM113109) and the Human Frontier Science Program (HFSP; grant number RGP0016/2019).

## DATA AVAILABILITY STATEMENT

The data that support the findings of this study are available from the corresponding author upon reasonable request.

## LIST OF ABBREVIATIONS

ac: anterior commissure
AIS: anterior intraencephalic sulcus
Ccer: commissure cerebelli
CeP: cerebellar plate
Ch: notochord
DIV: trochlear decussation
DVe: diencephalic ventricle
E: epiphysis
EmT: eminentia thalami
fp: floor plate
fr: fasciculus retroflexus
GM: gray matter
Ha: habenula
Hc: caudal hypothalamus
Hi: intermediate hypothalamus
Hr: rostral hypothalamus
Hy: hypophysis
III: oculomotor nerve
IL: inferior lobe of hypothalamus
IV: trochlear nerve
IX: glossopharyngeal nerve
le: lens
lfb: lateral forebrain bundle
LR: lateral recess of hypothalamic ventricle
lr: lateral rectus muscle
M: Mauthner neurons
M1: pretectal migrated area
M2: posterior tubercular migrated area
M3: EmT migrated area
M4: telencephalic migrated area
MHB: midbrain-hindbrain boundary
mlf: medial longitudinal fascicle
mou: mouth
MV: median ventricle of the mesencephalic ventricle
MVS: median ventricular sulcus of the mesencephalic ventricle
N: region of the nucleus of the medial longitudinal fascicle
NIII: oculomotor nucleus
NIn: interpeduncular nucleus
NIV: trochlear nucleus
NVa: rostral trigeminal motor neuron
NVII: facial motor neuron
NVp: caudal trigeminal motor neuron
OA: occipital arch
OB: olfactory bulb
oc: optic chiasma
OC: otic capsule
OE: olfactory epithelium
of: oval fossae
OG: octaval ganglion (VIII)
ON: olfactory nerve
on: optic nerve
ot: otolith
P: pallium
pc: posterior commissure
pf: pectoral fin
Po: preoptic region
poc: postoptic commissure
PoR: preoptic recess
PR: posterior recess of hypothalamic ventricle
Pr: pretectum
ptc: posterior tubercular commissure
PTd: dorsal part of posterior tuberculum
PTh: prethalamus
PTM: posterior tectal membrane
PTv: ventral part of posterior tuberculum
r: rhombomere
re: retina
RL: rhombic lip
RVe: rhombencephalic ventricle
S: subpallium
SC: spinal cord
Sd: dorsal subpallium
sr: superior rectus muscle
Sv: ventral subpallium
T: midbrain tegmentum
tc: tela choroidea
Tel: telencephalon
TeO: tectum opticum
TeVe: tectal ventricle
Th: thalamus
TL: torus longitudinalis
tpc: tract of posterior commissure
TS: torus semicircularis
TVe: telencephalic ventricle
V: trigeminal nerve
Va: valvula cerebeli
VI: abducens nerve
VII: facial nerve
WM: white matter
X: vagus nerve
Zli: zona limitans intrathalamica

## Notes

### Competing Interest Statement

The authors have declared no competing interest.

